# Structural snapshots of hyaluronan formation reveal principles of length control and secretion

**DOI:** 10.1101/2023.05.11.540447

**Authors:** Ireneusz Górniak, Zachery Stephens, Satchal K. Erramilli, Tomasz Gawda, Anthony A. Kossiakoff, Jochen Zimmer

## Abstract

Hyaluronan (HA) is an essential component of the vertebrate extracellular matrix. It is a heteropolysaccharide of alternating *N*-acetylglucosamine (GlcNAc) and glucuronic acid (GlcA) units reaching several megadaltons in healthy tissues. HA is synthesized and secreted in a coupled reaction by HA-synthase (HAS). Here, structural snapshots of HAS provide important insights into HA biosynthesis, from substrate recognition to HA elongation and translocation. We reveal a loop insertion mechanism for substrate binding, monitor the extension of a GlcNAc primer with GlcA, and capture the opening of a secretion channel that coordinates a nascent HA polymer. Further, we identify HA-interacting residues that control HA product lengths. Integrating structural and biochemical analyses, we propose a mechanism for HA length control based on finely tuned enzymatic processivity and catalytic rates.

## INTRODUCTION

Hyaluronan (HA) is an extracellular matrix polysaccharide of vertebrate tissues, with essential functions in osmoregulation, cell signaling and joint lubrication [1,2]. HA’s physiological functions are driven by its length and gel-forming properties [3]. Healthy tissues usually contain polymers of several megadaltons in size [2]. Inflammatory processes and osteoarthritis, however, associate with a decline in HA length to hundreds of kilodaltons [4,5].

HA consists of alternating N-acetylglucosamine (GlcNAc) and glucuronic acid (GlcA) units [6] and is synthesized and secreted by HA-synthase (HAS), a membrane-embedded processive glycosyltransferase (GT) [7–9]. HAS integrates several functions: first, binding its substrates (uridine diphosphate (UDP)-activated GlcA and GlcNAc); second, catalyzing glycosyl transfer; and third, translocating the nascent polymer across the plasma membrane. How these functions are integrated for HA biosynthesis is poorly understood.

While HA is ubiquitously expressed in vertebrates, the *Paramecium bursaria* chlorella virus (Cv) encodes a HAS similar to the vertebrate homologs [10]. Structural and functional analyses of Cv-HAS provided important insights into the enzyme’s architecture and how it initiates HA biosynthesis [11]. However, because the enzyme produces HA polymers substantially shorter than vertebrate HAS, as well as lacking a nascent HA chain in structural analyses, little was learned about how HAS coordinates the nascent HA polymer, translocates it between elongation steps, and regulates its size.

Biochemical and structural analyses of *Xenopus laevis* HAS isoform 1 (Xl-HAS-1, formerly DG42 [12,13]) address these critical questions. *In vitro*, Xl-HAS-1 produces polysaccharides similar in size to natively expressed HA. Cryogenic electron microscopy (cryoEM) analyses of the enzyme bound to a nascent HA chain reveal how it creates an HA translocation channel and coordinates the polysaccharide during secretion. Complementary cryoEM structures of Cv-HAS bound to the substrate UDP-GlcA and reaction products delineate the mechanism of alternating substrate polymerization and suggest a model for HA translocation. Further, site-directed mutagenesis of HA-coordinating residues identifies key interactions underlying HA length control.

## RESULTS

### Xl-HAS-1 synthesizes HA polymers in a megadalton size range

Xl-HAS-1 was expressed in *Spodoptera frugiperda* (Sf9) cells and purified using immobilized metal affinity (IMAC) and size-exclusion chromatography (SEC) in glyco-diosgenin (GDN) detergent (see Methods). Based on SEC and cryoEM analyses (see below), the catalytically active enzyme is monomeric (Fig. 1a,b).

**Figure 1.**
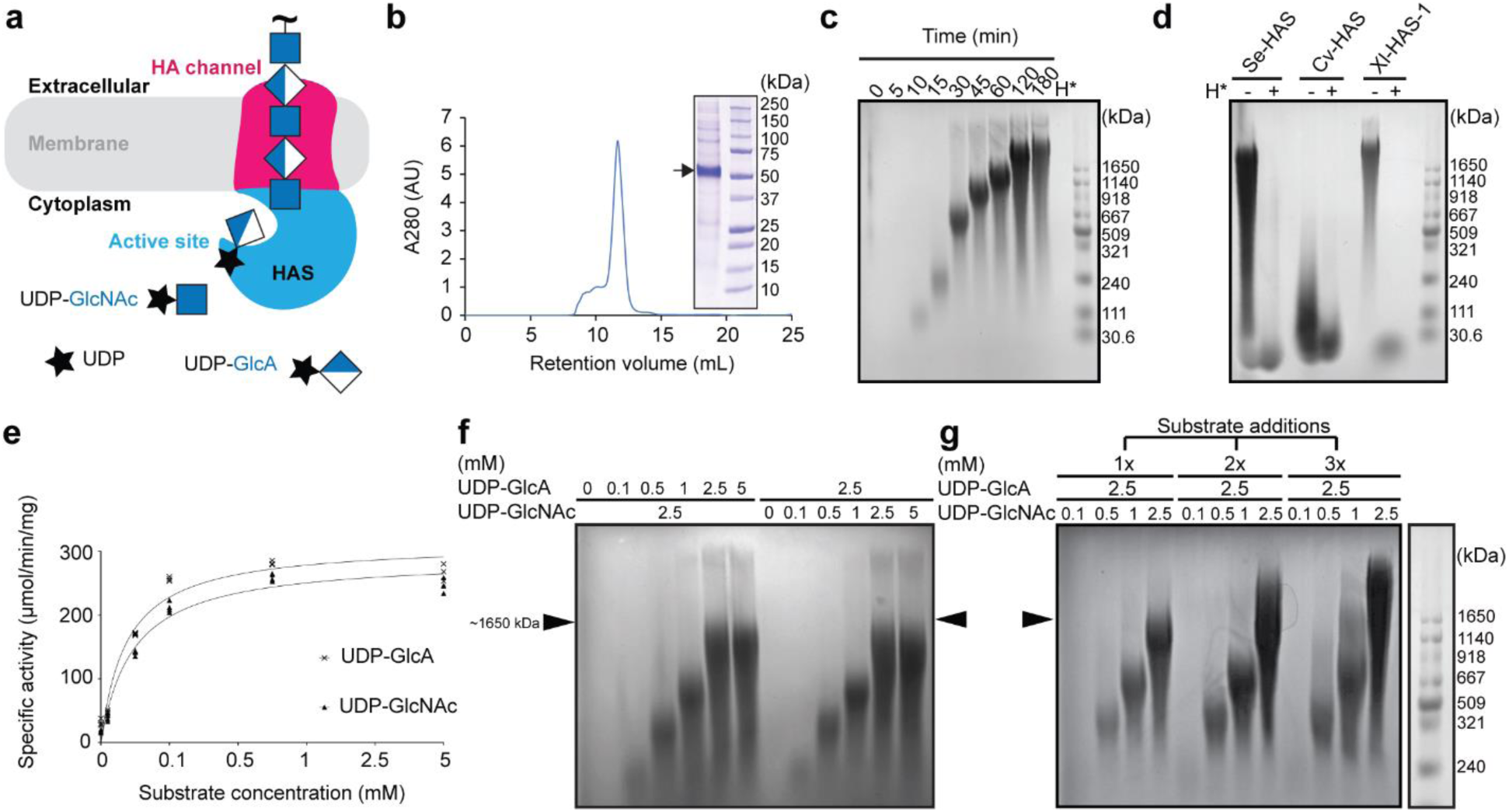
*In vitro* hyaluronan biosynthesis. (**a**) HAS couples HA synthesis with membrane translocation. (**b**) S200 size exclusion chromatogram of purified Xl-HAS-1. Inset: Coomassie-stained SDS-PAGE. (**c**) Time-course of *in vitro* HA biosynthesis monitored by agarose gel electrophoresis and Stains-all staining. H*: Hyaluronidase digestion. The molecular weight marker represents HA size standards. (**d**) Comparison of HA synthesized *in vitro* by *Streptococcus equisimilis* (Se), *Chlorella* virus (Cv) and *Xenopus laevis* (Xl) HAS. (**e**) Catalytic activity depending on substrate concentration. One substrate concentration was at 2.5 mM while the other was varied as indicated. Activity is measured by quantifying the released UDP in an enzyme coupled reaction. (**f**) The same as in panel (**e**) but by monitoring HA formation. Arrows correspond roughly to a 1.6 MDa HA marker. (**g**) Increasing HA molecular weight in the absence of product inhibition. A standard synthesis reaction was supplemented three times with 2.5 mM UDP-GlcA and the indicated amounts of UDP-GlcNAc while enzymatically converting UDP to UTP to prevent product inhibition.

HAS transfers the UDP-activated glycosyl unit (the donor sugar) to the non-reducing end of the nascent HA polymer (the acceptor sugar). This reaction generates UDP and HA as reaction products (Fig. 1a). Catalytic activity can be assessed by monitoring the release of UDP in an enzyme-coupled reaction, as previously described [8,11], or quantifying HA, either radiometrically or by dye staining following electrophoresis.

Xl-HAS-1 exhibits robust catalytic activity in the presence of magnesium cations, whereas its activity is reduced to about 30% in the presence of manganese. The synthesized polysaccharide is readily degraded by hyaluronidase, demonstrating that it is indeed authentic HA (Fig. S1a,b). Agarose gel electrophoresis of the synthesized HA followed by Stains-All staining reveals increasing polymer lengths over a ∼120 min synthesis reaction before stalling (Fig. 1c). Notably, at completion, Xl-HAS-1 synthesizes HA polymers of a consistent length that migrate above a 1.6 MDa HA marker. This material is comparable to HA *in vitro* synthesized by *Streptococcus equisimilis* (Se)-HAS (Fig. 1d). In contrast, Cv-HAS produces polydisperse HA of low molecular weight of up to 200 kDa.

Kinetic analyses were performed by titrating one substrate at a saturating concentration of the other and real-time quantification of the released UDP product (Fig. 1e). The results reveal Michaelis-Menten constants for UDP-GlcNAc and UDP-GlcA binding of about 470 and 370 µM, respectively, with an apparent maximum catalytic rate of approximately 20 substrate turnovers per minute per enzyme (Fig. S1g). Similar biosynthesis rates have recently been reported for cellulose synthase [14]. Monitoring the length distribution of the synthesized HA under these substrate conditions (one limiting, the other in excess) shows increasing HA lengths up to a substrate concentration of 2.5 mM (Fig. 1f).

*In vivo*, HA biosynthesis could stall due to substrate depletion, competitive UDP inhibition, and/or HA accumulation. To analyze the impact of sustained high substrate concentrations during HA biosynthesis, an *in vitro* synthesis reaction was supplemented with both substrates two, four, and six hours after initiation, while converting the accumulating UDP to non-inhibitory UTP (Fig. 1g and Fig. S1d-f). These conditions further extend the synthesized HA polymers to products substantially exceeding the 1.6 MDa marker. Extension is only observed when both substrates are supplied in excess (2.5 mM). If one substrate (UDP-GlcNAc) remains at a limiting concentration (0.1–1 mM), no increase in HA size is observed, perhaps due to premature HA release, followed by re-initiation.

### Architecture of Xl-HAS-1 in a resting state

We selected a non-inhibitory antibody Fab fragment to facilitate cryoEM structural analyses of Xl-HAS-1 (Fig. 2a,b and Fig. S1h,i). The purified Xl-HAS-1-Fab complex was either used directly for cryoEM grid preparation or incubated with substrates to obtain snapshots of biosynthetic intermediates (Fig. S2–4 and Table S1).

**Figure 2.**
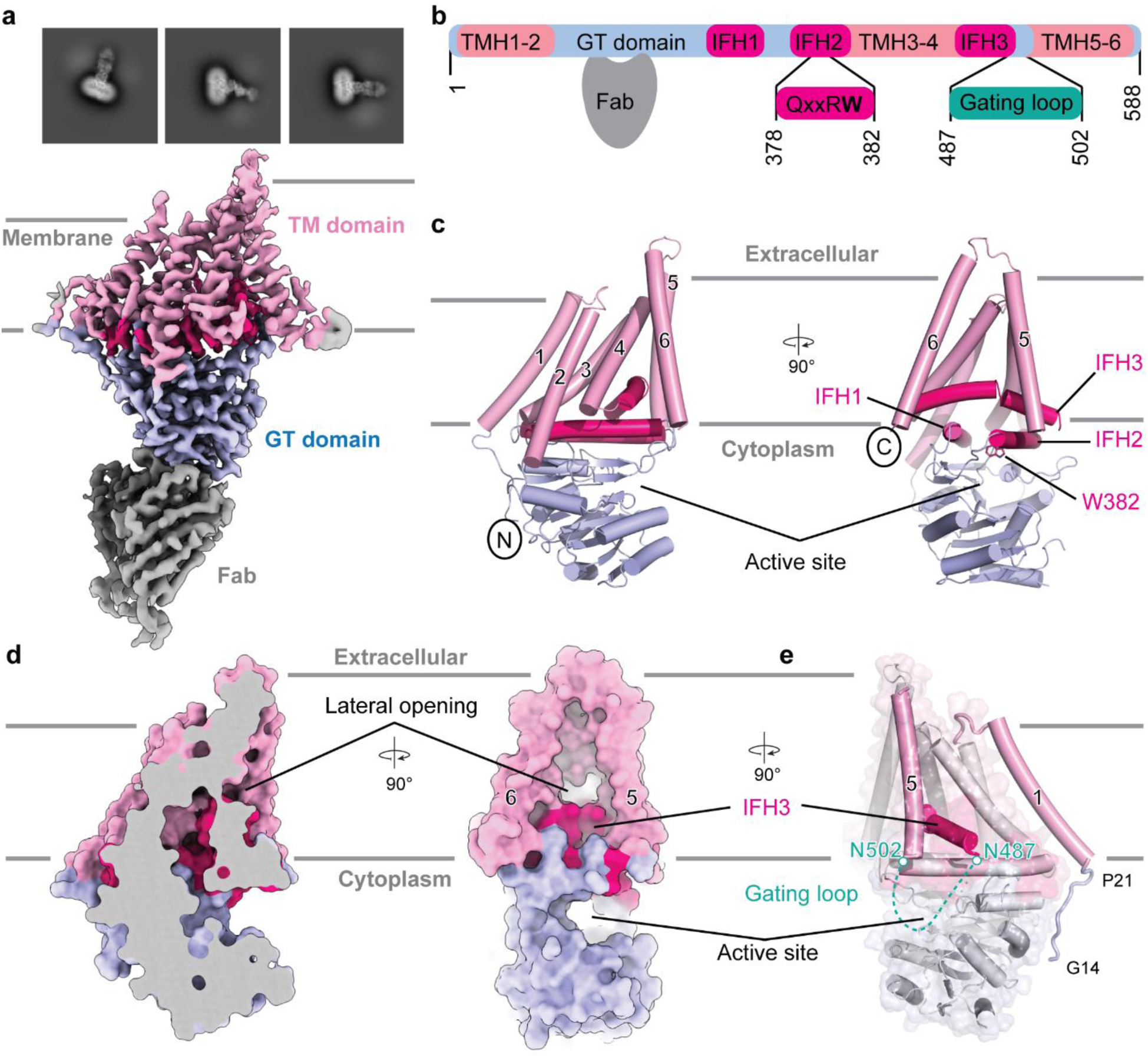
Structure of *Xenopus laevis* HAS-1. (**a)** Representative 2D class averages and electron potential map of Xl-HAS-1. The map was contoured at *σ* = 9 r.m.s.d. (**b)** Domain organization of Xl-HAS-1 and Fab binding site. (**c**) Architecture of Xl-HAS-1. The catalytic domain, amphipathic interface helices (IFH), and transmembrane regions are colored lightblue, hotpink, and pink, respectively. (**d)** Surface representation of Xl-HAS-1 indicating a curved channel with lateral opening. (**e**) Cartoon representation of Xl-HAS-1 indicating the positions of the N-terminal extension and the unresolved gating loop above the active site (dashed line).

Xl-HAS-1’s cytosolic GT domain interacts with two N-terminal and four C-terminal transmembrane helices (TMHs) as well as three cytosolic interface helices (IFH1-3) (Fig. 2b,c). The IFHs surround the entrance to a curved channel that ends with a lateral portal halfway across the membrane, formed by IFH3 and TMHs 5 and 6 (Fig. 2d). The function of the curved channel is unknown as it is not used for HA translocation, except for its cytosolic entrance (see below). IFH2 contains the conserved QxxRW motif (Fig. S5) of which Trp382 is pivotal for positioning the acceptor sugar right at the entrance to the TM channel.

The Xl-HAS-1 architecture is consistent with an AlphaFold predicted structure of human HAS-1 and 2 (Fig. S6a) and the previously determined Cv-HAS structure [11], with the exception that TMH1 is clearly resolved in the Xl-HAS-1 maps (Fig. 2e and Fig. S6b). This curved helix rests against the cytosolic connection of IFH2 and TMH3 (Fig. 2e).

Vertebrate HASs contain a conserved WGTSGRR/K motif in a cytosolic ‘gating loop’ connecting IFH3 with TMH5. A similar motif in a loop close to the active site is found in chitin and cellulose synthases [11,15–17] (Fig. S6d-f). Although the gating loop is unresolved in the apo Xl-HAS-1 structure, its flanking residues Asn487 and Asn502 at the C-terminus of IFH3 and beginning of TMH5, respectively, position it right above the catalytic pocket (Fig. 2e) where it could facilitate substrate binding [11].

### Structural snapshots of HA biosynthesis intermediates

To stabilize an HA-associated translocation intermediate, we performed an *in vitro* HA synthesis reaction in the presence of hyaluronidase to trim polymers emerging from HAS’ TM channel. The purified reaction intermediates were either used directly for cryoEM grid preparation (approach #1) or supplemented with additional UDP-GlcNAc to control the HA register (approach #2).

### HAS forms an electropositive HA translocation channel

CryoEM maps of HA-bound Xl-HAS-1 obtained from approach #1 reveal continuous non-proteinaceous density running from the catalytic pocket to its extracellular surface through an electropositive channel (Fig. 3a,b, and Fig. S3b). The density accommodates nine glycosyl units, denoted 1–9 starting at the non-reducing end located at the acceptor site. The HA map at the first and fifth glycosyl position strongly suggests the presence of GlcNAc units with well-resolved acetamido groups (Fig. S3c). This assignment is consistent with previous analyses of Cv-HAS demonstrating stable coordination of GlcNAc, but not GlcA, at the acceptor position established by Trp382 of the QxxR**W** motif [11] (Fig. 2c and 3a). The remainder of the HA polymer was modeled with the carboxylate and acetamido groups of a disaccharide repeat unit pointing roughly in the same direction, as observed in crystallographic structures and by *in silico* modeling (Fig. S6c) [18] [19].

**Figure 3.**
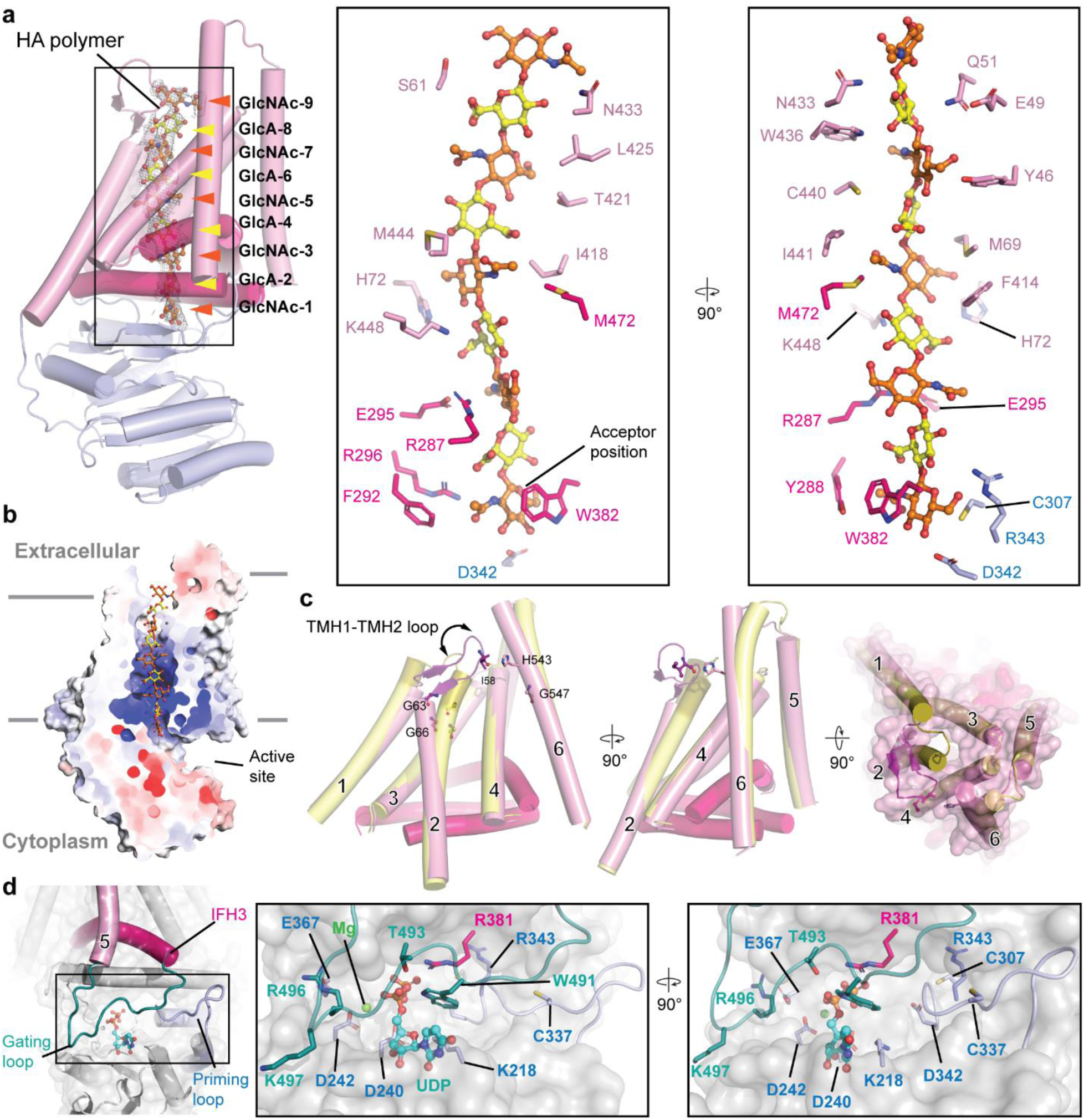
A HA-HAS translocation intermediate. (**a)** Left panel: Xl-HAS-1 structure with a bound HA nonasaccharide. The cryoEM map is shown as a black mesh contoured at *σ* = 6 r.m.s.d. Right panel: Stick representation of residues coordinating HA inside Xl-HAS-1 TM channel. (**b**) Electrostatic surface potential of Xl-HAS-1 calculated in PyMOL using the APBS plugin [41,43]. The nascent HA polymer is shown as ball- and-sticks. (**c**) Gating transitions upon HA coordination. Shown is a superimposition of the TM regions only in the presence (pink) and absence (yellow) of a translocating HA polymer. (**d**) Gating loop insertion into the catalytic pocket. Conserved residues contacting UDP are shown as sticks. UDP is shown as ball-and-sticks colored cyan for carbon atoms.

At the acceptor position near the active site, the first GlcNAc unit (GlcNAc-1) sits inside a collar of invariant residues, including Trp382 stacking against the sugar ring, Asp342 in hydrogen bonding distance to GlcNAc-1’s C3 hydroxyl group, and Arg343 contacting the C6 hydroxyl of the terminal GlcNAc unit (Fig. 3a). Asp342 and Arg343 belong to the conserved GD**DR** motif found in HA synthases (Fig. S5). Furthermore, Arg296 interacts with GlcNAc-1’s acetamido group, and the sulfhydryl group of Cys307 is located within 4.5 Å to the sugar’s pyranose ring. The aromatic residues Tyr288 and Phe292 complete a pocket that accommodates the hydrophobic part of GlcNAc-1’s acetamido group. The carboxylate of the following GlcA-2 is stabilized by Arg296.

While the first two glycosyl units of HA enter the channel in a flat conformation, the polymer rotates by about 90° around the GlcA-2 – GlcNAc-3 glycosidic linkage (Fig. 3a and Fig. S6c). The rotation is induced by a salt bridge extending into the channel volume, formed between the conserved Arg287 and Glu295 side chains. His72 and Lys448 form an electropositive pocket that accommodates the carboxylate of GlcA-4. The orientation of the next disaccharide unit (GlcNAc-5 and GlcA-6) is well-defined by the GlcNAc-5 density (Fig. S3c). It is rotated by ∼180° relative to the preceding GlcNAc-GlcA pair and sits in a Met-rich ring of hydrophobic residues, including Met69, Phe414, Ile418, Ile441, Met444, and Met472 (Fig. 3a).

In the second half of the TM channel, HA is surrounded by moderately conserved hydrophilic and hydrophobic residues, including Tyr46, Glu49, Gln51, Ser61, Thr421, Leu425, Asn433, Trp436, and Cys440. These residues mediate long-distance interactions with the hydroxyl groups of the glycosyl units (Fig. 3a).

### Bending of TMHs opens the HA translocation channel

HA translocation requires conformational changes that create a continuous TM channel. Compared to the resting conformation described above, the N-terminal half of TMH2 near the extracellular water-lipid interface bends away from TMH4 (Fig. 3c). Bending occurs around a conserved GLYG motif (residues 63 to 66) that places two Gly residues at the interface with TMH4. Furthermore, the last and first helical turns of TMH1 and TMH2, respectively, unwind to form a short β-hairpin with residues of the TMH1–2 loop (Fig. 3c). This loop lids the extracellular channel exit in the closed conformation. Upon channel opening, the loop flips towards the membrane and rotates by approximately 90° to run roughly parallel to the membrane surface. Additionally, the N-terminal half of TMH6 bends by approximately 10° towards TMH2, around the conserved Gly547 (Fig. 3c and Fig. S5). In the new position, His543 is in hydrogen bonding distance to the backbone carbonyl oxygen of Ile58.

### HAS’ gating loop coordinates the nucleotide at the active site

The second approach to trap a Xl-HAS-1-HA translocation intermediate resulted in a state in which the enzyme is bound to UDP with a closed TM channel (Fig. 3d and Fig. S4). UDP is coordinated via conserved motifs of the GT domain, as previously described for Cv-HAS as well as other processive and non-processive GTs of the GT-A fold [11,15,16,20–22]. Additionally, Xl-HAS-1’s gating loop, connecting IFH3 with TMH5 (residues 488 to 501) and containing the conserved WGTSGRK/R sequence, is resolved in this map (Fig. S4 and 6). The loop inserts into the catalytic pocket to coordinate the nucleotide (Fig. 3d). In this conformation, Trp491 forms a cation-π interaction with Arg381 of the Qxx**R**W motif in IFH2, which in turn forms a salt bridge with the nucleotide’s diphosphate group. The indole ring of Trp491 runs approximately perpendicular to the uracil moiety, such that its Nɛ is within hydrogen bonding distance of UDP’s α-phosphate. This phosphate group is also contacted by the hydroxyl group of the following Thr493. The side chain of Arg496, the penultimate residue of the WGTSG**R**K/R motif, is incompletely resolved, but likely interacts with the equally conserved Glu367 of the GT domain to stabilize the gating loop in the inserted conformation.

### The gating loop is necessary for HA biosynthesis

Site-directed mutagenesis was used to reveal the gating loop’s functional importance in two model systems, Xl-HAS-1 as well as Cv-HAS. Enzymatic activities were monitored with purified enzyme for Xl-HAS-1 or in inverted membrane vesicles (IMVs) for Cv-HAS (Fig. 4a-d).

**Figure 4.**
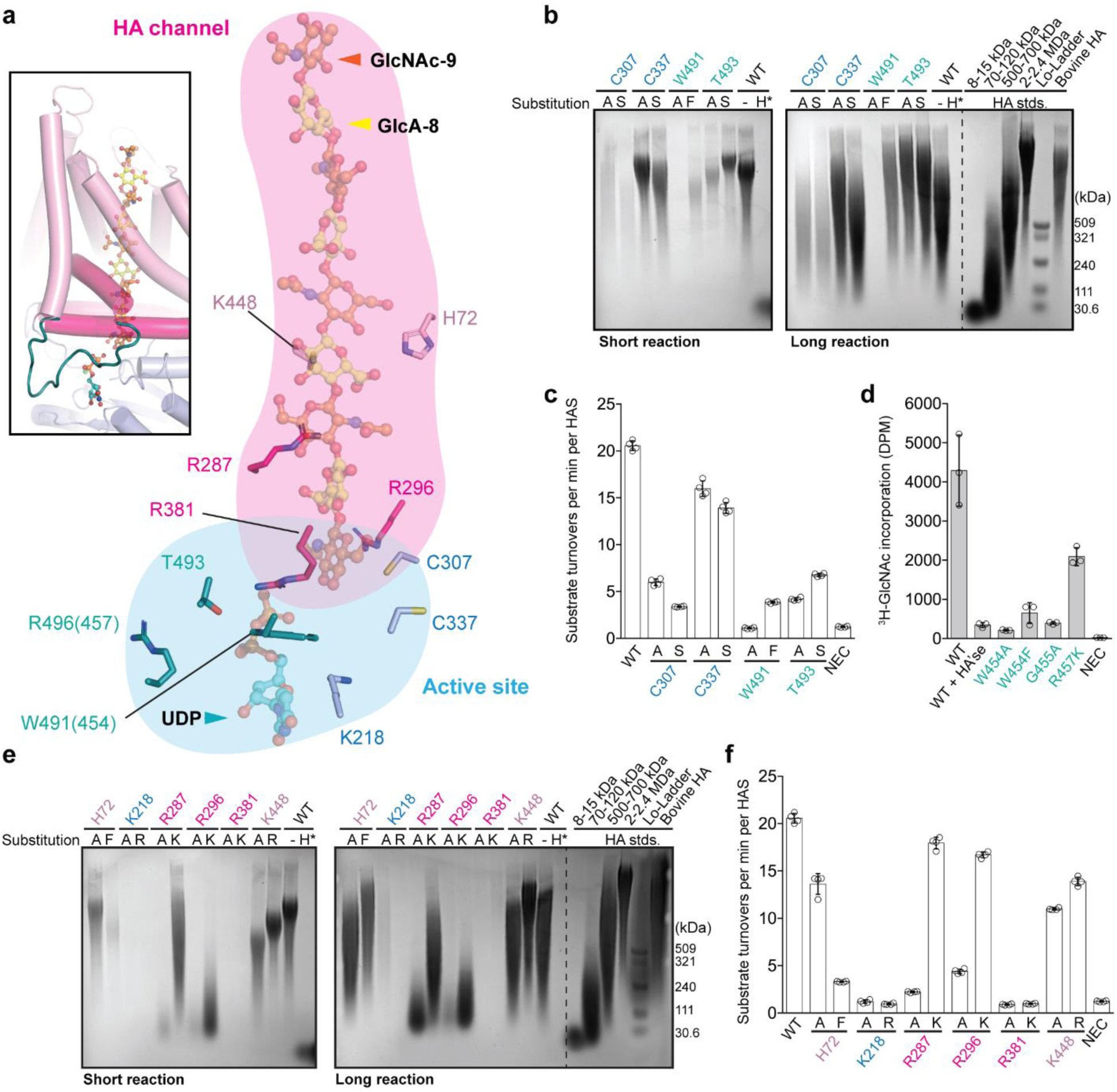
Modulation of HA length control. (**a**) Overview of mutagenized residues forming the active site (residues colored blue and deepteal correspond to the GT domain and gating loop, respectively) or the HA channel (residues colored pink and hotpink for the TM domain and IFH, respectively). Residue numbers in brackets refer to the Cv-HAS sequence. (**b**) Mutagenesis of active-site-lining residues. *In vitro* HA biosynthesis by the indicated Xl-HAS-1 mutants for 1 h or overnight (short and long reactions, respectively). WT: Wild-type. HA standards (HA stds.) are shown to the right of the gel depicting the overnight synthesis results. Lo: Low molecular weight. Agarose gels were stained with Stains-all. (**c**) Catalytic rates of the mutants shown in (**b**). Rates report the substrate turnovers per min and HAS enzyme and were determined by quantifying UDP release in real time during HA synthesis. (**d**) Catalytic activity of gating loop mutants of Cv-HAS. Activity was determined by quantifying ^3^H-labeled HA by scintillation counting [8] and are reported relative to the wild-type activity. DPM: Disintegrations per minute. (**e**) Mutagenesis of channel-lining residues. *In vitro* HA biosynthesis by the indicated mutants, similar to panel (**b**). (**f**) Similar to panel (**c**) but for channel-lining residues. The experiments were performed at least in triplicate (*n*=3 or 4) and error bars represent standard deviation from the means.

For Xl-HAS-1, replacing Trp491 of the **W**GTSGRK/R motif with Ala abolishes HA biosynthesis during a 60-min synthesis reaction (denoted ‘short reaction’), while its substitution with Phe results in a lower molecular weight product (Fig. 4a-c). An overnight synthesis reaction (‘long reaction’) with this mutant produces HA polymers similar to the wild-type enzyme. Replacing the following Thr493 (WG**T**SGRK/R) with Ala or Ser reduces the catalytic rate to about 20–25% of that of the wild-type (Fig. 4b,c). In a short synthesis reaction, the T493A and T493S mutants generate HA lengths comparable to and exceeding the wild-type product, respectively (Fig. 4b). These product size differences are less apparent after a long synthesis reaction. For all reactions, additional shorter HA polymers accumulate overnight, likely due to substrate depletion and premature termination of biosynthesis.

Similar results were obtained for the viral enzyme by quantifying HA using scintillation counting (Fig. 4d). Here, the gating loop contains a WGTRG sequence. Replacing the motif’s Trp residue (Trp454) with Ala or Phe is incompatible with function and so is the substitution of the following Gly455 with Ala (W**G**TRG). Replacing the Arg residue (Arg457) with Lys reduces Cv-HAS’s activity by ∼60% relative to the wild-type.

### HA coordination modulates polymer length

Processive HA biosynthesis requires sustained interactions of HAS with HA between elongation steps. To test how HA coordination affects the HA length distribution, we substituted several HA- coordinating residues.

Xl-HAS-1’s active site contains two conserved cysteines. Cys307 belongs to the ‘switch loop’ at the back of the active site where it coordinates the acceptor sugar (Fig. 3a and 4a). Replacing Cys307 with Ala or Ser leads to the production of low amounts of polydisperse HA (Fig. 4b,c). The second Cys, Cys337, is part of a priming loop located to one side of the catalytic pocket. This residue can be replaced with Ala or Ser, resulting in HA products similar to wild-type, with a slight length reduction for the C337S mutant (Fig. 4b). The corresponding residue in Cv-HAS is Cys297 (see below) and its replacement with Ala renders Cv-HAS inactive [11].

Also at the active site, replacing the conserved Lys218 with Ala or Arg abolishes catalytic activity in short and long synthesis reactions (Fig. 4e,f). Similarly, Arg381 of the Qxx**R**W motif, located in IFH2 right above the catalytic pocket, cannot be replaced with either Ala or Lys. This residue interacts with the nucleotide’s pyrophosphate group in UDP- and substrate-bound states (see below).

Inside the TM channel, the polymer’s second and third glycosyl units (GlcA-2 and GlcNAc-3) are near Arg287 of IFH1. This residue induces HA rotation past GlcA-2 due to forming a salt bridge with Glu295 (Fig. 3a and 4a). Substituting Arg287 with Ala generates only low-molecular-weight HA in short and long synthesis reactions. Replacing it with Lys, however, results in a mutant with roughly wild-type activity producing polydisperse HA (Fig. 4e,f). Similarly, Arg296 of the acceptor binding pocket, also located in IFH1, is critical for function. Neither an Ala nor Lys substitution yields HA products comparable to the wild-type enzyme. Overnight, the R296K mutant produces short HA fragments as also observed for the R287A construct. Lastly, catalytically active enzymes are obtained when K448, about 4.5 Å away from the carboxylate of HA’s GlcA-4, is replaced with Ala or Arg. The mutants produce HA polymers of defined lengths, although at reduced catalytic rates, with the K448A mutant being the slowest. Strikingly, in a long reaction, the variants produce polymers equivalent to or exceeding the HA length obtained from the wild-type enzyme (Fig. 4e,f). Lastly, replacing His72 at the interface of TMH2 and TMH3 with Ala shows no HA length variation but an overall decreased product yield (Fig. 4a,e,f). Replacing it with Phe, the corresponding residue in Cv-HAS, reduces the enzyme’s catalytic activity substantially in a short synthesis reaction.

### Substrate binding repositions the GlcNAc primer

Upon priming with a GlcNAc monosaccharide [11], HAS attaches GlcA to the C3 hydroxyl group of the primer to form the GlcA-GlcNAc disaccharide repeat unit. To gain insights into this step, we took advantage of the high-quality cryoEM maps routinely obtained for Cv-HAS, facilitated by two camelid nanobodies [11]. The catalytically inactive Cv-HAS D302N mutant [11,23] in complex with two nanobodies was incubated with GlcNAc and Mn^2+^:UDP-GlcA prior to grid preparation (Fig. S7).

The GlcNAc primer is well resolved in the cryoEM map and occupies the same acceptor position as previously reported (Fig. 5a) [11]. In the presence of the UDP-GlcA substrate, however, the primer rotates by approximately 60° to position its acetamido group away from the active site (Fig. 5a). This rotation positions the primer’s C3 hydroxyl group closer to the base catalyst Asp302 (replaced with Asn in this construct).

**Figure 5.**
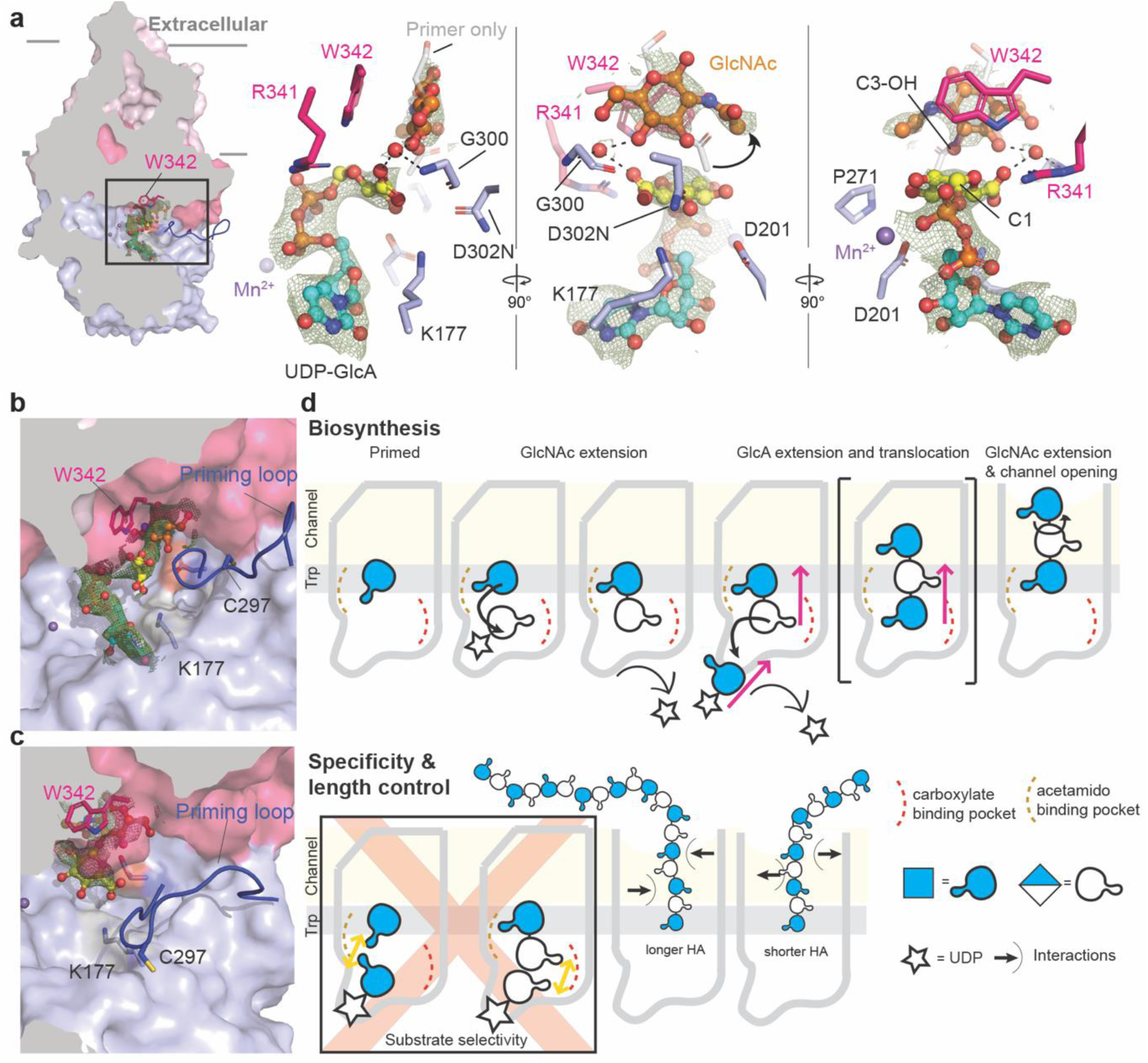
Substrate binding and primer extension by Cv-HAS. (**a)** UDP-GlcA bound to GlcNAc-primed Cv-HAS. Ligands are shown as ball-and-sticks and the corresponding EM map as a light-green mesh contoured at *σ*=5 r.m.s.d. The priming sugar is colored orange and the UDP-GlcA substrate yellow and cyan for the carbon atoms of the donor sugar and uracil moiety, respectively. The primer in the absence of a donor substrate is shown as gray and red sticks (from PDB: 7SPA). (**b)** and (**c**) Extension of the GlcNAc primer by GlcA. (**b**) Disaccharide- and UDP-bound Cv-HAS. (**c)** Disaccharide-bound Cv-HAS. Cv-HAS is shown as a surface with selected residues of the active site as sticks for orientation. The EM maps for the ligands are shown as a green mesh contoured at σ=4.5 r.m.s.d and *σ*=4 r.m.s.d. for the UDP-bound, and nucleotide-free states. (**d)** Model of HA biosynthesis. Top: The priming GlcNAc is positioned at the acceptor site (gray bar labeled ‘Trp’). UDP-GlcA binding leads to glycosyl transfer and formation of a disaccharide. UDP-GlcNAc binding induces disaccharide translocation and glycosyl transfer to form a trisaccharide transiently positioned with GlcA at the acceptor site. HA may rotate inside the channel. Bottom, left: Consecutive incorporation of the same glycosyl unit is prevented due to overlapping coordination sites of the acetamido and carboxylate substituents. Right: The coordination of HA by HAS and catalytic rates modulate HA length distribution.

Cv-HAS binds its substrates in similar binding poses, with UDP being coordinated by two manganese ions, as previously described [11] (Fig. 5a). The donor GlcA sugar sits roughly underneath Trp342 of the QxxR**W** motif. Its C6 carboxylate is incompletely resolved, as often observed in electron potential maps. The carboxylate occupies a pocket formed by the C-terminal end of the priming loop (residues 298 to 300), the following helix that begins with the invariant GDDR motif (residues 300-303), as well as the priming GlcNAc sugar (Fig. 5a). Further, Arg341 of the Qxx**R**W motif interacts with the donor’s carboxylate. Additional long distance (up to 4.5 Å) interactions exist between the GlcA donor and Asp201 (via its C3 hydroxyl), as well as Lys177 and the introduced Asn302 (D302N) (via its C4 hydroxyl). The distance between the acceptor’s C3 hydroxyl group and the donor’s C1 carbon atom is long (about 5.2 Å), indicating that additional conformational changes are necessary for glycosyl transfer. In the substrate-bound states, Cv- HAS’ priming loop with the conserved Cys297 at its tip is retracted from the active site (Fig. 5a,b), as described previously for UDP and UDP-GlcNAc bound states [11].

### HA translocates in disaccharide steps

We also attempted to visualize intermediate HA translocation states of viral HAS. To this end, the wild-type Cv-HAS-nanobody complex was incubated with both substrates prior to EM analysis. This resulted in Cv-HAS structures bound to a HA disaccharide, with GlcNAc at the acceptor site and GlcA extending into the catalytic pocket. In this position, GlcA is flexible and incompletely resolved in the EM map. One disaccharide bound state also contained UDP, corresponding to a state directly after glycosyl transfer, prior to UDP release (Fig. 5b,c and Fig. S8).

In the UDP-bound state, the newly added GlcA unit remains in close proximity to UDP’s β-phosphate (Fig. 5b). Because the UDP and HA disaccharide densities are not clearly separated, we cannot exclude the presence of an overlapping minor second state in which Cv-HAS is bound to UDP-GlcA and the GlcNAc monosaccharide primer (Fig. 5b).

In the absence of UDP, the priming loop inserts into the catalytic pocket, and the HA disaccharide continues to extend into the catalytic pocket with its GlcA moiety (Fig. 5c). This suggests that the extended HA does not translocate spontaneously into the TM channel, even after priming loop insertion into the active site. We note that the carboxylate group of GlcA, either when extending the priming GlcNAc unit or as part of the donor sugar, occupies the same pocket of the catalytic site (Fig. S9).

## DISCUSSION

HA, cellulose and chitin synthases are multitasking enzymes that secrete high molecular weight polysaccharides. How the enzymes control product lengths and thus physical properties is largely unresolved. The narrow size distribution of the Xl-HAS-1-synthesized HA suggests minimal termination and re-initiation events. The size distribution broadens significantly during prolonged *in vitro* synthesis reactions, likely due to substrate depletion and/or product inhibition. Thus, *in vivo*, substrate availability and local UDP concentrations could modulate HA length [24].

Processive HA biosynthesis requires a stable HA-HAS association. Accordingly, altering HA coordination inside the secretion channel profoundly affects the HA length distribution. Two Arg-to-Lys substitutions within Xl-HAS-1’s IFH1 (R287K and R296K) abolish length control or lead to early termination of HA biosynthesis. These residues at the entrance to the TM channel are likely critical in stabilizing the nascent chain for elongation. Additionally, replacing K448, about halfway across the channel, with Ala or Arg substantially reduces the enzyme’s catalytic rate, leading to shorter yet discrete product lengths. Notably, mutations reducing Xl-HAS-1’s catalytic rate while not changing its processivity give rise to higher molecular weight HA under extended synthesis conditions. *In vivo*, similar effects may be achieved by limiting substrate availability and/or post-translational modifications [25,26]. Thus, differences in HA length associated with HAS isoforms could result from different metabolic states of the expressing cells and tissues. The shape and electropositive character of Xl-HAS-1’s TM channel contrast the flat, acidic channel formed by cellulose synthase [20,27]. Because the channel’s gate is near the extracellular water–lipid interface, we estimate channel opening is induced by HA polymers exceeding 3-4 glycosyl units (Fig. 5d). In this case, the nascent chain and the channel’s central hydrophobic ring likely prevent water flux across the membrane, similar to other polysaccharide secretion systems [28].

Our new viral HAS structures demonstrate that, upon extending a GlcNAc primer with GlcA, the newly added sugar protrudes into the catalytic pocket. In this position, the terminal GlcA would overlap with the sugar moiety of the next donor substrate. Therefore, we propose that binding of a new substrate molecule induces HA translocation, due to a steric clash (Fig. 5d). We further assume the simultaneous formation of a new linkage between GlcA and the incoming donor sugar (GlcNAc). This would transiently position a trisaccharide with GlcA at the acceptor site and GlcNAc extending into the catalytic pocket. MD simulations show that this register is unstable, leading to spontaneous sliding of the HA polymer into the channel by one glycosyl unit, thereby again stably positioning a GlcNAc unit at the acceptor site [11] (Fig. 5d).

This model assumes that HA enters the TM channel with the carboxylate and acetamido groups of a disaccharide repeat unit pointing in opposite but always the same directions, respectively (Fig. 5d). Past the second sugar position, the channel architecture and dimension may allow HA to rotate into an energetically more favorable conformations with the carboxylate and acetamido groups of a disaccharide repeat unit pointing roughly in the same direction (Fig. S6c) [19]. In particular, the Glu295-Arg287 salt bridge protruding into the channel may initiate this rotation (Fig. 3a).

Lastly, comparing the structures of GlcNAc-primed HAS bound to its substrates suggests a mechanism for alternating substrate polymerization (Fig. 5d). Accordingly, extending a terminal glycosyl unit with an identical sugar is prevented due to overlapping binding pockets of their main substituents (Fig. S9). At the acceptor and donor positions, the acetamido group of GlcNAc or carboxylate of GlcA would occupy the same binding pocket. Similar clashes are prevented during alternating substrate polymerization because the GlcNAc’s and GlcA’s substituents are localized on opposite sides of the glucopyranose ring (Fig. 5d).

Our structural and functional analyses provide a molecular framework for engineering polysaccharide biosynthesis for a plethora of biomedical, agricultural, and engineering purposes.

## METHODS

### Xl-HAS-1 expression

The synthetic gene encoding N-terminally dodeca-histidine-tagged Xl-HAS-1 (Uniprot ID: P13563) was cloned into the pACEBac-1 vector using BamHI and HindIII restriction sites. Xl-HAS-1 mutants were generated using QuickChange and Q5 SDM methods [29,30]. Cloning was confirmed by restriction analyses and DNA sequencing. Baculoviruses harboring each target gene were prepared as described [27]. Briefly, the Xl-HAS-1 containing pACEBac-1 plasmid was transformed into *E. coli* DH10MultiBac cells. Bacmids for the wild type and all Xl-HAS-1 mutants were purified from 3 white colonies and transfected into Sf9 cells at 1×10^6^ cells/mL using the FuGene reagent (Promega). Cells were maintained in ESF921 medium (Expression Systems) at 27 °C with mild shaking. The baculovirus was amplified to generate P2 virus stock, which was used at 1.5% culture volume to infect Sf9 cells at a density of 3×10^6^ cells/mL. Cells grown in 1 L media bottles (0.45 L per bottle) were pelleted by centrifugation 48 h post-infection and resuspended in 40 mL of Buffer A (40 mM TrisHCl pH 7.8, 150 mM NaCl, 10 % glycerol, 5 mM MgCl_2_) per 0.9 L of cell culture. The harvested cells were flash-frozen and stored at −80 °C for subsequent use.

### Cv-HAS expression

Expression of Cv-HAS was performed as described [11]. Briefly, electrocompetent *E. coli* C43 cells were transformed with a pET28a-Cv-HAS expression vector. LB medium supplemented with 50 µg/mL kanamycin was inoculated with the transformed C43 cells and grown overnight in an orbital shaker at 37 °C, 220 rpm. The overnight culture was used to inoculate 4 L of terrific broth supplemented with 1xM salts (25 mM Na_2_HPO_4_, 25 mM KH_2_PO_4_, 50 mM NH_4_Cl, 5 mM Na_2_SO_4_) [31] and 50 µg/mL kanamycin. Cultures were grown at 30 °C with 220 rpm shaking to OD600=0.8, then cooled to 20 °C for 1 hour. To induce expression, isopropyl-β-D-thiogalactopyranoside was added to a final concentration of 100 µg/mL. After 18 hours, cells were harvested by centrifugation at 5000 rpm in a JLA-8.1000 rotor for 20 minutes. Cell pellets were flash-frozen in liquid nitrogen before storage at −80 °C.

### Xl-HAS-1 purification

All preparation steps were carried out at 4 °C unless stated otherwise. To purify Xl-HAS-1, typically 0.9 L-culture worth of cell suspension was thawed and diluted to 200 mL using Buffer A supplemented with 5 mM β-ME, 1 mM phenylmethylsulfonyl fluoride (PMSF), 10 mM imidazole, 1% β-dodecyl maltoside (DDM) and 0.2% cholesteryl hemisuccinate (CHS). Cells were lysed using a tissue homogenizer and rocked for one hour at 4 °C, followed by ultracentrifugation for 30 minutes. The cleared lysate was mixed with 10 mL of 50% Ni-NTA (ThermoFisher) resin suspension equilibrated in Buffer A and subjected to batch binding for an hour. After that, the slurry was poured into a glass gravity flow column (Kimble), the flow-through was discarded, and the resin was washed three times with 50 mL of Buffer A supplemented with 0.03% GDN (wash 1) and 1 M NaCl (wash 2) or 20 mM imidazole (wash 3). Xl-HAS-1 was eluted using Elution Buffer (EB) consisting of 25 mM Tris-HCl pH 7.8, 150 mM NaCl, 10% glycerol, 350 mM imidazole, 0.03% GDN in two steps. First, 15 mL of EB was added followed by ∼ 5 min incubation, draining, and addition of another 15 mL of EB and draining. The eluted sample was concentrated using 50 kDa MWCO Amicon centrifugal concentrator (Millipore) to <1 mL for size exclusion chromatography (SEC) using a Superdex200 column (GE healthcare) equilibrated in Gel Filtration Buffer (GFB) consisting of 20 mM Tris-HCl pH 7.8, 150 mM NaCl and 0.02% GDN. The target peak fractions were pooled and concentrated as necessary for subsequent experiments.

### Cv-HAS purification

Purification of Cv-HAS was performed as described [11]. All steps were carried out at 4 °C. Harvested C43 cells were resuspended in RB (20 mM Tris pH 7.5, 100 mM NaCl, 10% glycerol, 0.5 mM TCEP-HCl), dounce homogenized and mixed with 1 mg/mL egg white lysozyme for 1 hour. Cells were lysed by three passages through a microfluidizer at 18,000 psi, 2 mM PMSF was added after the first passage. Unbroken cells and debris were cleared by centrifugation at 12,500 rpm in a JA-20 rotor. Supernatant from the JA-20 spin was collected to harvest membranes by ultracentrifugation in a Ti-45 rotor at 42,000 rpm for 2 hours. The crude membrane fraction was dounce homogenized in SB (20 mM Tris pH 7.5, 300 mM NaCl, 40 mM imidazole, 10% glycerol, 0.5 mM TCEP, 1% DDM, 0.1% CHS, 1 mM PMSF) and mixed for one hour by gentle inversion. Insoluble material was pelleted by ultracentrifugation at 42,000 RPM in a Ti-45 rotor for 30 minutes. The supernatant was mixed in batch with 5 mL Ni-NTA resin (ThermoFisher), equilibrated in SB, for one hour. Flow-through material was collected by gravity, after which the nickel resin was washed with 20 bed volumes WB1 (20 mM Tris pH 7.5, 1 M NaCl, 10% glycerol, 40 mM imidazole, 0.02% DDM, 0.002% CHS, 0.5 mM TCEP) and 20 volumes WB2 (20 mM Tris pH 7.5, 300 mM NaCl, 10% glycerol, 80 mM imidazole, 0.02% DDM, 0.002% CHS, 0.5 mM TCEP). Cv-HAS was eluted in 5 volumes EB (20 mM Tris pH 7.5, 300 mM NaCl, 10% glycerol, 320 mM imidazole, 0.02% DDM, 0.002% CHS, 0.5 mM TCEP). The eluate was concentrated and injected on an S200 Increase 10/300 GL column (Cytiva) equilibrated in GFB1 (20 mM Tris pH 7.5, 100 mM NaCl, 0.5 mM TCEP, 0.02% DDM and 0.002% CHS). Fractions containing Cv-HAS were pooled and concentrated for subsequent reconstitution.

### Cv-HAS inverted membrane vesicle preparation

To prepare Cv-HAS-bearing inverted membrane vesicles (IMVs), 2 L-culture worth of cell suspension was thawed in a water bath for 10 minutes at 30 °C. 5 mM β-mercaptoethanol (β-ME) and 1 mM PMSF were added and the cells were lysed in a microfluidizer. Unbroken cells were pelleted by centrifugation at 5000 ×*g* for 15 minutes in a JA-20 rotor, followed by layering the cleared lysate onto 40 mL of 2 M sucrose cushion and ultracentrifugation in Ti-45 rotor (Beckman) at 42,000 rpm for 2 hours. The brown IMV ring was collected, diluted to 60 mL using Buffer A (20 mM Tris-HCl pH 7.5, 100 mM NaCl, 10% glycerol) and subjected to another round of ultracentrifugation for 1 hour. The pelleted IMVs were resuspended in 2 mL of Buffer A, aliquoted, flash-frozen, and stored at −80 °C for subsequent use in activity assays.

### Quantification of HAS activity by scintillation counting

All reagents for biochemical analyses were supplied by Sigma, unless stated otherwise. To quantify the bulk Xl-HAS-1 activity, radioactive labeling assay was employed (*8*). Reaction buffer consisted of 25 mM TrisHCl pH 7.8, 150 mM NaCl, 0.02% GDN, 20 mM MgCl_2_, 5 mM UDP-GlcA, 5 mM UDP-GlcNAc and 0.01 µCi/μL of [^3^H]-UDP-GlcNAc (PerkinElmer). Reactions were carried out at 37 °C for 3 hours, and at 1 µM HAS concentration. Negative controls were carried out by either omitting the enzyme (background radioactivity) or digesting the synthesized polymers with bovine hyaluronidase (MP Biochemicals, 1 mg/mL, 10 minutes at 30 °C). The synthesized HA polymer was purified by descending paper chromatography and quantified using liquid scintillation counting, as described [8].

Quantification of Cv-HAS activity followed a similar workflow. Protein concentrations in CvHAS IMVs were normalized as described previously [14]. Reaction buffer consisted of 40 mM TrisHCl pH 7.5, 75 mM NaCl, 20 mM MnCl_2_, 0.5 mM TCEP, 5 mM UDP-GlcNAc, 5 mM UDP-GlcA and 0.01 μCi/μL [^3^H]-UDP-GlcNAc. Reactions were carried out at 30 °C for 2 hours. HA digests were performed by adding 2% DDM and 75 U hyaluronidase to the reaction and incubating for an additional 10 minutes at 30 °C. Reactions were quenched with 3% SDS, and subsequent product was quantified using scintillation counting.

### Electrophoretic HA size determination

To assess the size of HA synthesized by Xl-HAS-1, similar reaction conditions were applied to the ones described above, except the radioactive tracer was omitted. Synthesis reactions were mixed with SDS-PAGE loading dye and applied to a 1% agarose gel (Ultra-pure agarose, Invitrogen) casted in an Owl B2 system (ThermoFisher). All gels were subjected to electrophoresis at 100 V for 2 hours at room temperature to achieve comparable separation for each run. After the run, the gel was equilibrated in 50% ethanol for 1.5 h and subjected to staining in 0.05% Stains-All (Sigma) in 50% ethanol overnight under light-protection. Post-staining background was reduced by soaking the gel in 20% ethanol for 3-7 days in the dark. The migration of the synthesized HA species was compared to HA standards from *Streptococcus equi* (Sigma), as well as enzymatically generated ladders: HA LoLadder and HA HiLadder (Hyalose).

### Substrate turnover rate quantification

UDP release during HA synthesis was quantified using an enzyme-coupled assay as previously described [8,11]. Depletion of NADH was monitored at 340 nm every 60 s for 3 hours at 37 °C in a SpectraMax instrument. The raw data was processed in MS Excel. The rate of NADH depletion was converted to µmoles of UDP released using a UDP standardized plot (Fig. S1c) for Michaelis-Menten constant determination using GraphPad Prism. For substrate turnover rate quantification of Xl-HAS-1 mutants, µmoles of released UDP were converted to the number of corresponding substrate turnovers per molecule of Xl-HAS-1, per minute. All experiments were performed in quadruplicate and error bars represent the standard deviation from the means.

### Xl-HAS-1 reconstitution and identification of specific Fabs

For Fab selection, Xl-HAS-1 was reconstituted into *E. coli* total lipid nanodiscs [32] using chemically biotinylated MSP1D1. The purified enzyme was mixed with MSP and sodium cholate-solubilized lipids at 40 µM final concentration in 1 mL final volume according to 1:4:80 molar ratio of HAS:MSP:lipids. Detergent removal was initiated 1 hour after mixing all ingredients by adding 200 mg of BioBeads (BioRad) and mixing at 4 °C. After an hour, another batch of BioBeads was added followed by mixing overnight. The following morning (after ∼12 hours), the reconstitution mixture was transferred to a fresh tube and the last batch of BioBeads was added followed by mixing for one hour and SEC using Superdex200 column equilibrated with GFB lacking detergent.

Phage selection was performed as previously described [33,34]. In the first round of selection 400 nM of Xl-HAS-1-loaded nanodiscs diluted in the selection buffer (20 mM HEPES pH 7.5, 150 mM NaCl, 1% BSA) was immobilized on streptavidin paramagnetic beads (Promega). Beads were washed three times in the selection buffer, with 5 mM d-desthiobiotin added during the first wash to block nonspecific binding. Fab phage library E [35] resuspended in selection buffer was added to the beads and incubated for 1 hour with gentle shaking. The beads were washed three times in the selection buffer and then transferred to log-phase *E. coli* XL1-Blue cells. Phages were amplified overnight in 2xYT medium with ampicillin (100 µg/ml) and M13-KO7 helper phage (10^9^ pfu/mL). Four additional rounds of selection were performed with decreasing target concentration (200 nM, 100 nM, 50 nM, 25 nM) using a KingFisher magnetic beads handler (ThermoFisher). In every subsequent round amplified phage pool from the previous round was used as the input. Prior to being used for selection, each phage pool was precleared by incubation with 100 µl of streptavidin paramagnetic beads. Additionally, 2 µM of non-biotinylated MSP1D1 nanodiscs were present in the selection buffer to reduce the presence of non-specific binders during rounds two to five. In these rounds selection buffer supplemented with 1% Fos-choline-12 was used to release the target and bound phages from the nanodiscs. Cells infected after the last round were plated on LB agar with ampicillin (100 µg/ml) and phagemids from individual clones were sequenced at the University of Chicago Comprehensive Cancer Center Sequencing Facility to identify unique binders. Single-point phage ELISA was used to validate specificity of unique binders as described previously [34]. Fabs were expressed and purified as described [34] and used for activity assays and SEC co-elution experiments with Xl-HAS-1. The strongest non-inhibitory binders were chosen for cryoEM trails and one of those yielded well-structured projections of the HAS-Fab complex.

### Cv-HAS nanodisc reconstitution and nanobody complex formation

Nanobodies were expressed and purified as described earlier [11] and stored in aliquots at −80 °C for further use. Fractions from the initial SEC containing Cv-HAS were pooled, concentrated and combined with MSP1E3D1 and *E. coli* Total Lipid Extract (solubilized in 60 mM DDM) at a 1:3:30 ratio. When reconstituting Cv-HAS in a ternary complex with camelid nanobodies, the initial reconstitution mixture was supplemented with a 3-fold molar excess of appropriate Nb composition (3:3:1 Nb872:Nb881:CvHAS for the GlcNAc primed, UDP-GlcA state and 3:3:1 Nb872:Nb886:CvHAS for disaccharide associated states). The reconstitution was mixed by inversion for 30 minutes before adding BioBeads (BioRad) to about 1/3 of the total volume. After 30 minutes, an additional volume of BioBeads was added, and the reconstitution mixture was allowed to incubate overnight. The next day, another volume of BioBeads was added. After 30 minutes, the mixture was cleared of BioBeads, as well as lipid and protein aggregates by passing through a 0.2 µm cellulose acetate spin filter. The filtered mixture was re-injected on an S200 SEC column equilibrated in GFB2 (20 mM Tris pH 7.5, 100 mM NaCl, 2 mM MnCl_2_, 0.5 mM TCEP). Cv-HAS nanodisc fractions were screened by SDS-PAGE for co-eluting complex components.

### Xl-HAS-1 cryoEM sample preparation

GDN-solubilized Xl-HAS-1 was used for cryoEM experiments. The purified enzyme was mixed with Fab at a 1:4 molar ratio and incubated overnight at 4 °C, followed by SEC using Superose 6 column (Cytiva) equilibrated with GFB2 containing 0.01% GDN.

Initial attempts to prepare cryoEM grids of the purified Xl-HAS-1-Fab complex in the presence of substrates failed to capture a HAS–HA intermediate due to rapid HA accumulation in the sample hampering grid vitrification process. To generate a HA-associated sample, substrates (UDP-GlcA and UDP-GlcNAc, 2.5 mM each, Sigma) and 20 mM MgCl_2_ were included during the overnight Fab incubation, followed by 1 hour hyaluronidase (1 mg/mL) treatment and SEC. Post-SEC, the sample was concentrated to 8 mg/mL using a 100 kDa MWCO Amicon ultrafiltration membrane. To generate the GlcNAc-terminated, HA- and UDP-bound sample, 2.5 mM UDP-GlcNAc and 2.5 mM MgCl_2_ were added after SEC and incubated on ice for 1 hour prior to cryoEM grid vitrification. 4 µL of Xl-HAS-1 sample was applied onto the C-flat 1.2/1.3 grid, glow-discharged for 45 seconds in the presence of 1 drop of amylamine. Grids were blotted for 4 seconds at a blot force of 4, at 4 °C and 100% humidity and plunge-frozen in liquid ethane using Vitrobot Mark IV (FEI).

### Cv-HAS cryoEM sample preparation

For the GlcNAc-primed, UDP-GlcA-bound state of Cv-HAS, 5 mM GlcNAc was included in the initial nanodisc reconstitution mixture. Purified Cv-HAS nanodiscs in complex with Nb881 and Nb872 were concentrated to 3.5 mg/mL, then further supplemented with 10 mM MnCl_2_, 5 mM GlcNAc, and 5 mM UDP-GlcA, and incubated on ice for 10 minutes. Quantifoil R1.2/1.3 300 mesh grids were glow-discharged for 45 seconds with 1 drop of amylamine. A sample volume of 2.5 µl was applied to each grid at 4 °C, 100% humidity, blotted for 12 seconds with a blot force of 4 and plunge frozen in liquid ethane using Vitrobot.

For the Cv-HAS sample with observed HA oligosaccharide density, purified Cv-HAS in complex with Nb886 and Nb872 was concentrated to 7.0 mg/mL. The purified complex was diluted 2-fold with a reaction mixture containing 10 mM UDP-GlcNAc, 10 mM UDP-GlcA and 40 mM MnCl_2_. After mixing, the sample was incubated at room temperature for 15 minutes before applying 2.5 µL to glow discharged QF grids. Blotting parameters were adjusted to 6 seconds at a blot force of 6.

### CryoEM data collection and processing

All cryoEM datasets were collected on a Titan Krios equipped with a K3/GIF detector (Gatan) at the Molecular Electron Microscopy Core (University of Virginia School of Medicine). Forty-five-frame movies were recorded in counting mode at 81,000x nominal magnification, −2.0 to −1.0 µm target defocus, and 50 e^-^/Å^2^ total dose.

All datasets were processed in cryoSPARC [36]. Raw movies were subjected to patch motion correction and patch contrast transfer function (CTF) estimation. Particles were automatically selected by blob picker (Xl-HAS-1 HA-bound and UDP-bound datasets) or template picker (all other datasets), used to generate ab initio models and subsequently sorted by iterative cycles of 2D classification and heterogenous refinement. Due to high particle density, duplicates were removed from the blob-picked particle stack in cryoSPARC. To separate HASs bound to their respective ligands, 3D variability and 3D classification approaches were used. The final volumes were refined using non-uniform and local refinement to generate high-resolution maps with estimated resolutions from 3.0–3.6 Å (Fig. S2–4 and S7–8).

### Model building

To generate the Xl-HAS-1 model, the Alphafold2 [37] prediction was docked into the EM map using Chimera [38] and the model was iteratively real-space refined in Coot [39] and Phenix [40].

We were able to model most of Xl-HAS-1 residues with the exception of the unresolved loop of the GT domain (residues 172-193), the N-terminal (residues 1-14) and C-terminal (residues 569- 588) extensions, as well as the Gating loop (residues 487-502, except for the UDP-bound Xl-HAS- 1). For HA-bound Xl-HAS-1, HA register within the channel was assigned using an auto-sharpened map (MAP2) generated in Phenix, which showed improved density for the first and fifth sugar of the polymer (Fig. S3c). The final model was refined against MAP1 (Fig. S3).

Substrate- and HA oligo-bound Cv-HAS models were generated by docking in a proper ligand and real-space refining previously published Cv-HAS model (PDB ID 7SP6).

All model figures were prepared using PyMOL [41] or ChimeraX [42].

## Acknowledgments

We thank Kelly Dryden and Michael Purdy of the Molecular Electron Microscopy Center at the University of Virginia for support and are indebted to Ruoya Ho for help with cryoEM data collection. We thank Finn Maloney, Jeremi Kuklewicz and Louis Wilson for critical comments on the manuscript. We are also grateful to Paul DeAngelis for advice on HA detection and providing HA ladders. The project was in part funded by NIH grant R35GM144130 (to J.Z.) and R01GM117372 (to A.A.K.). I.G. is a recipient of the Boehringer Ingelheim Fonds Fellowship. J.Z. is an Investigator of the Howard Hughes Medical Institute. This article is subject to HHMI’s Open Access to Publications policy. HHMI lab heads have previously granted a nonexclusive CC BY 4.0 license to the public and a sublicensable license to HHMI in their research articles. Pursuant to those licenses, the author-accepted manuscript of this article can be made freely available under a CC BY 4.0 license immediately upon publication.

## Author contributions

J.Z., I.G. and Z.S. designed the experiments. I.G. performed all structural and functional analyses of Xl-HAS-1. Z.S. performed all structural and functional analyses of Cv-HAS. S.K.E., T.G. and A.A.K. selected Fab antibodies against Xl-HAS-1. All authors evaluated and interpreted the data. I.G., Z.S. and J.Z. wrote the manuscript and all authors edited it.

## SUPPLEMENTARY FIGURE TITLES AND LEGENDS

**Figure S1.**
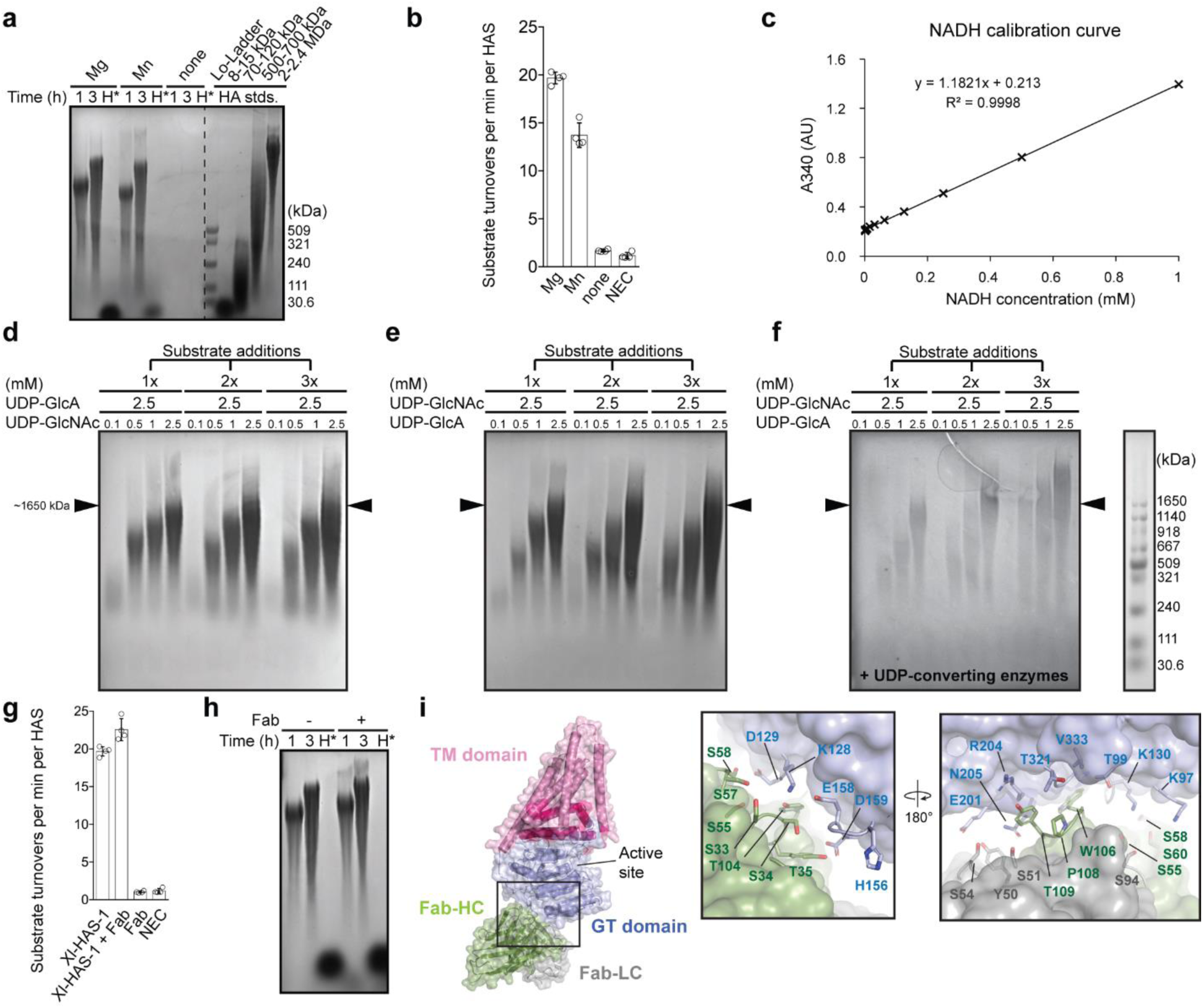
*In vitro* catalytic activity of Xl-HAS-1. (**a**) Agarose gel assay showing HA synthesis is dependent on Mg^2+^, while Mn^2+^ reduces HA production slightly. H* – hyaluronidase digestion. (**b**) Same as panel A but quantified using radioactive GlcNAc incorporation. NEC- no enzyme control. This experiment was performed in triplicate (n=3) and error bars represent standard deviations from the means. (**c**) NADH calibration curve used for substrate turnover rate calculations. (**d**) Substrate replenishing in the absence of UDP-converting enzymes (Lactate dehydrogenase – LDH and pyruvate kinase - PK). UDP-GlcA=2.5 mM const. and UDP-GlcNAc is varied for this assay. (**e**) Same as panel (**d**), except that UDP-GlcA is varied and UPD-GlcNAc is kept constant. (**f**) Substrate replenishing with LDH/PK present (UDP-GlcNAc=2.5 mM const), reverse to what is shown in Fig. 1g. Arrows indicate maximum HA extension without UDP removal and at 2.5 mM of both substrates, corresponding roughly to 1.6 MDa HA marker. (**g**) Substrate turnover rates in the presence and absence of the Fab. This experiment was performed in quadruplicates (n=4) and error bars represent standard deviations from the means. (**h)** HA synthesis in the presence and absence of the Fab. (**i**) HAS-Fab interface. Fab binds the GT domain at a position roughly opposite to the active site.

**Figure S2.**
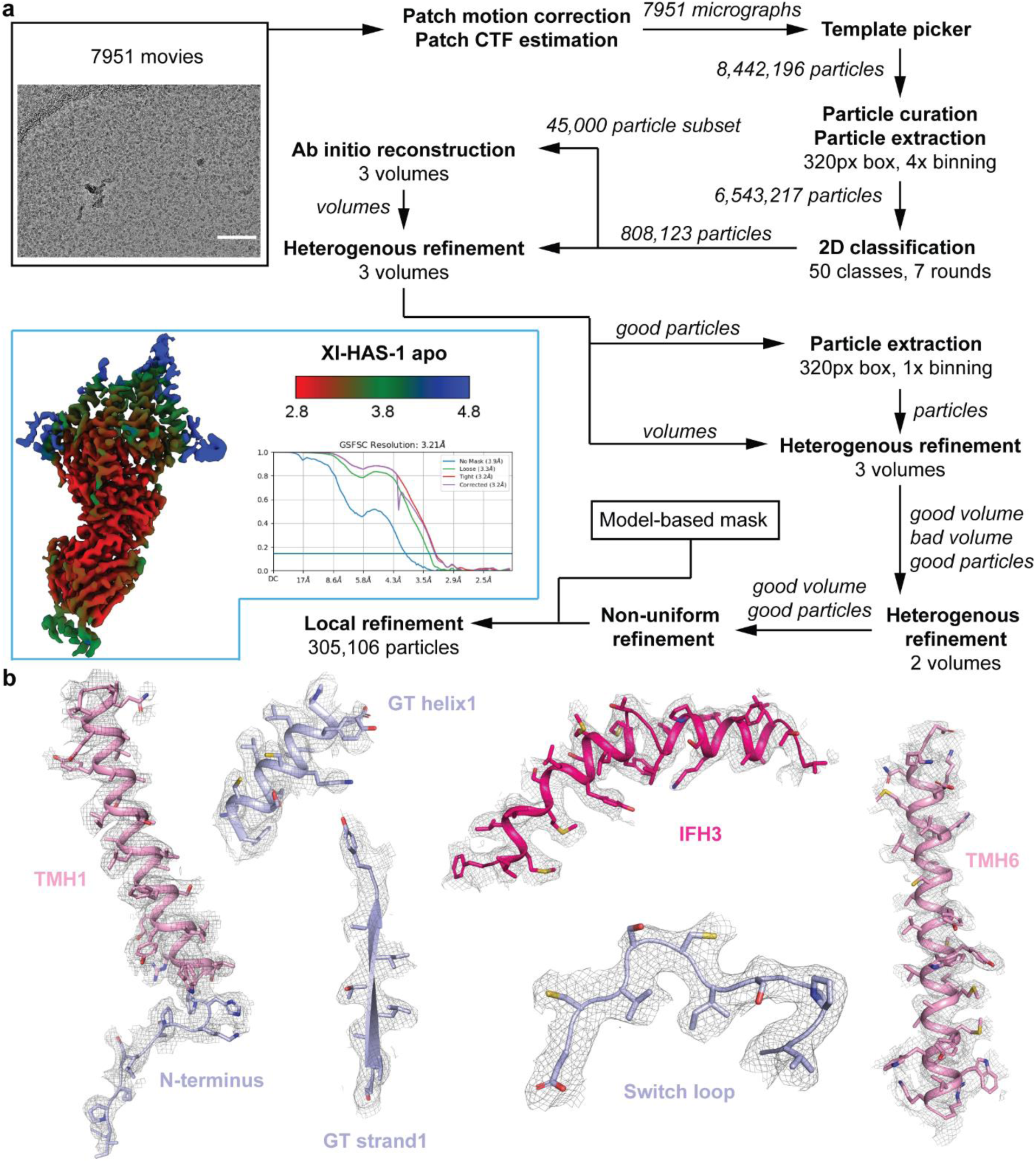
CryoEM data processing for apo Xl-HAS-1. **(a)** CryoSPARC workflow for apo Xl-HAS-1. **(b)** Representative regions showing map quality. Scale bar on the micrograph corresponds to 200 nm. The map was contoured at *σ*=6 r.m.s.d.

**Figure S3.**
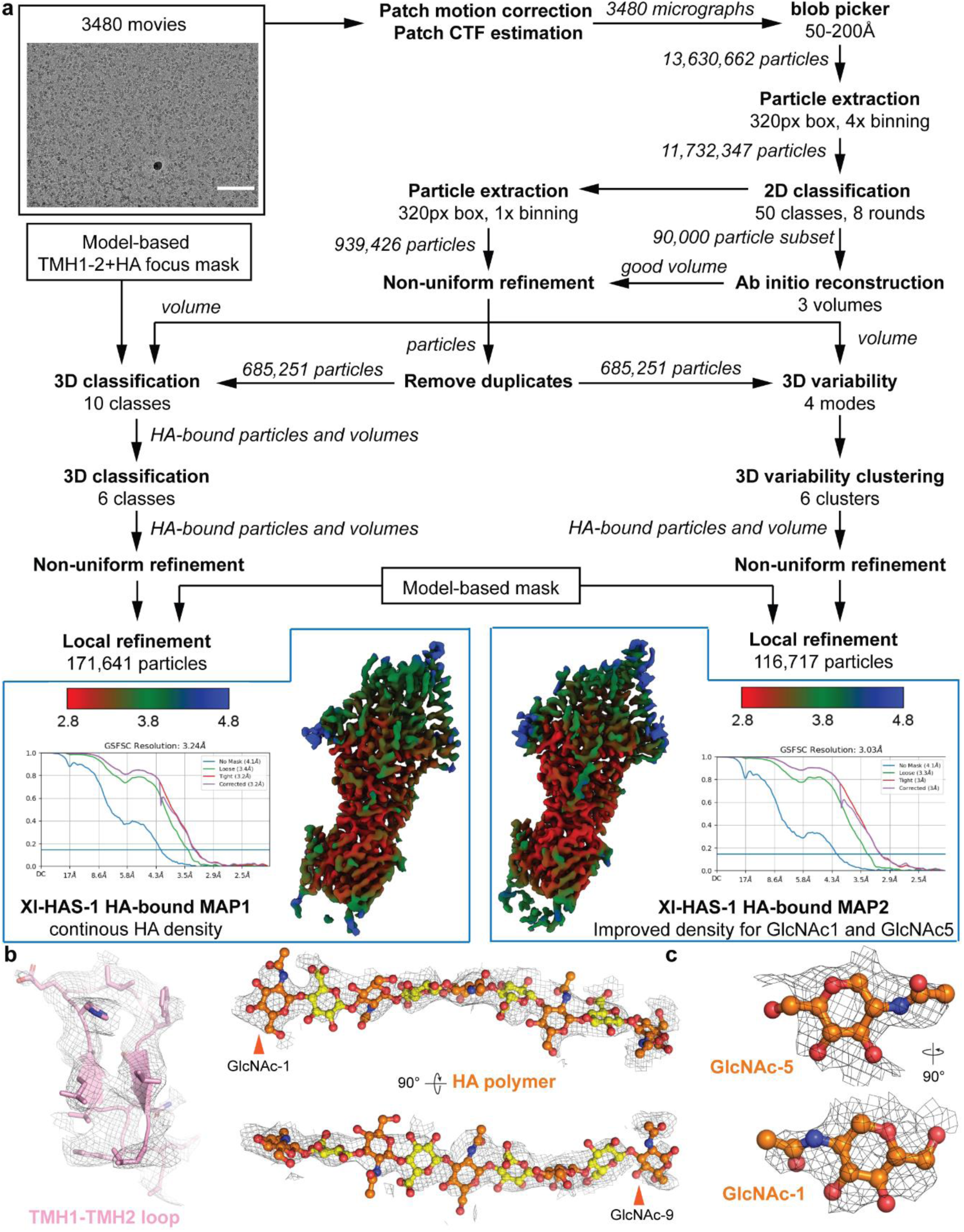
CryoEM data processing for HA-bound Xl-HAS-1. **(a)** CryoSPARC workflow for HA-bound Xl-HAS-1. **(b)** Map1 quality for relevant parts of the model. **c**, Map quality for GlcNAc1 and GlcNAc5. Scale bar on the micrograph corresponds to 200 nm. Maps were contoured at *σ*=6 r.m.s.d.

**Figure S4.**
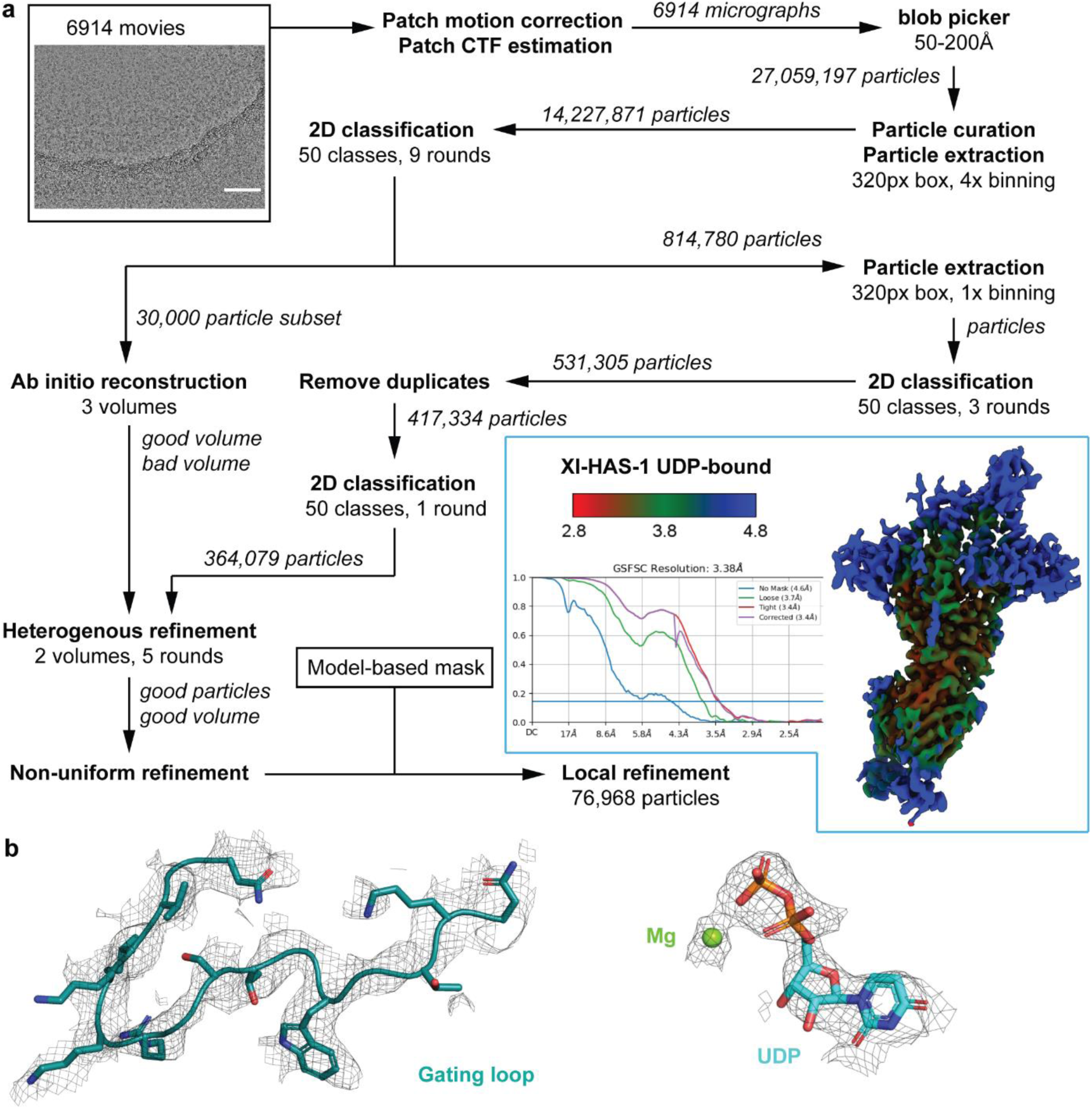
CryoEM data processing for UDP-bound Xl-HAS-1. **(a)** CryoSPARC workflow for UDP-bound Xl-HAS-1. **(b)** Map quality for the gating loop and UDP:Mg. Scale bar on the micrograph corresponds to 200 nm. The map was contoured at *σ*=7 r.m.s.d.

**Figure S5.**
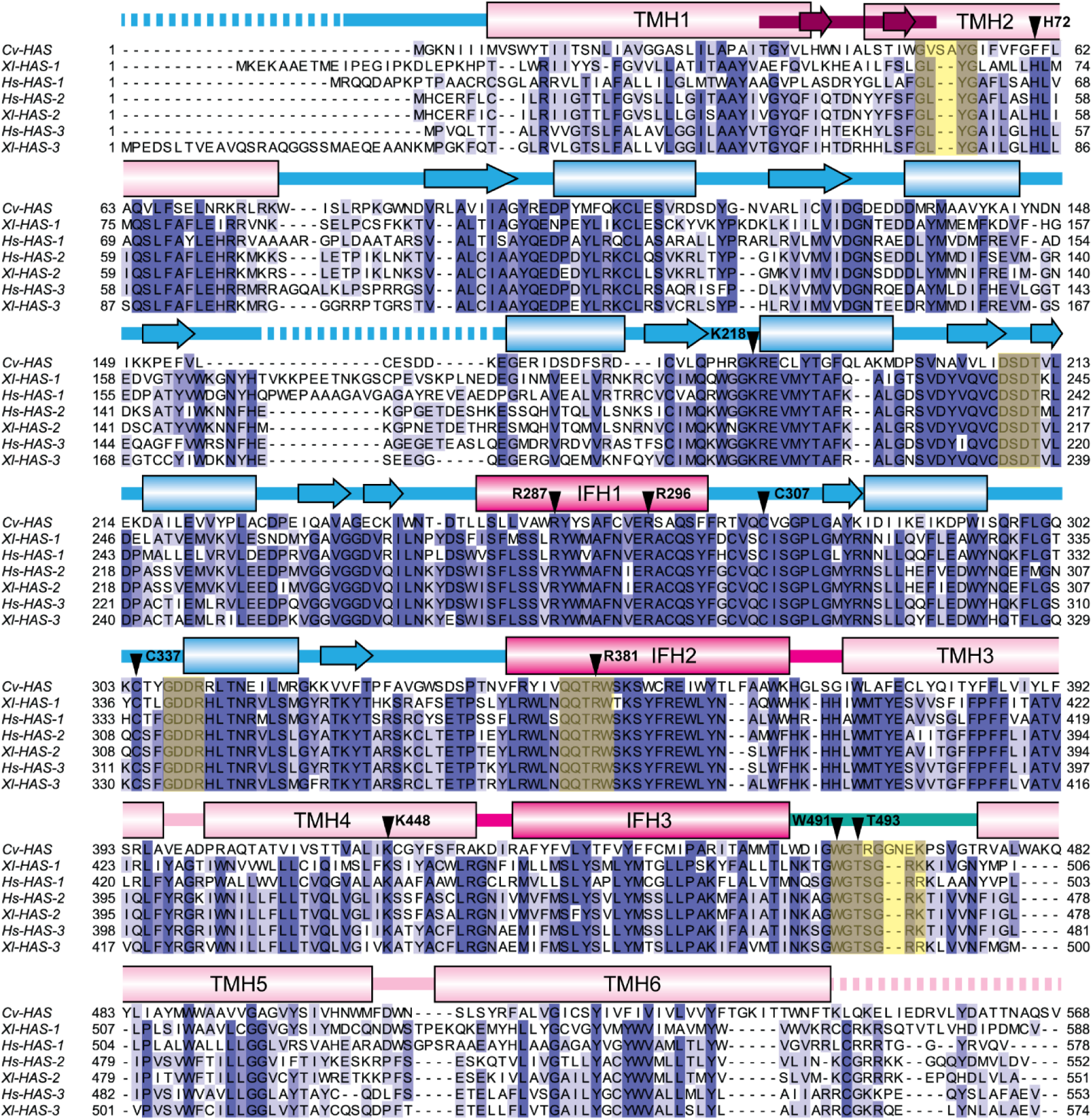
HAS sequence alignment. HAS sequence alignments (MUSCLE) [44] showing crucial conserved motifs (GLYG, DxDT, GDDR, QxxRW and WGTSGRK/R, highlighted in yellow), as well as the conserved residues used for mutagenesis studies (indicated with arrowheads). The bar on top of the sequences is colored according to structural domains of Xl-HAS-1 (pink: TM domain, purple: TMH1-TMH2 loop, hotpink: IF domain, lightblue: GT domain, deepteal: Gating loop). Cylinders indicate α-helices, arrows β-sheets, lines loops, while dashed lines correspond to unstructured regions.

**Figure S6.**
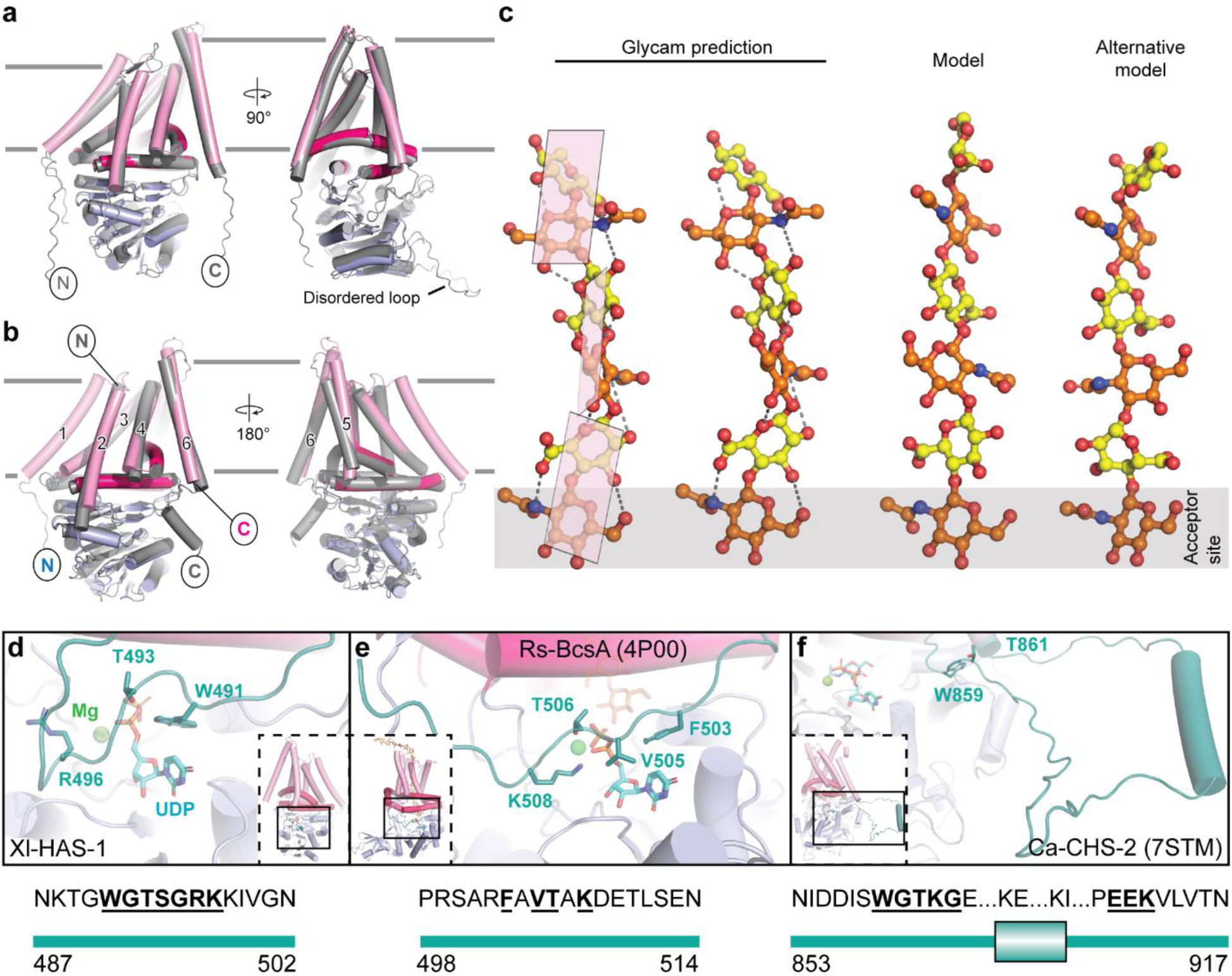
HA modeling and gating loop comparison. **(a)** Superimposition of Xl-HAS-1 apo (colored) and Hs-HAS-1 AlphaFold 2 prediction (gray). (**b**) Superimposition of Xl-HAS-1 apo (colored) and Cv-HAS (gray, PDB: 7SP6). (**c)** Conformation of HA. Left: Glycam predicted structure of an HA hexasaccharide (glycam.org). Middle: Conformation of HA as modeled in the Xl-HAS-1 complex. Right: Alternative but likely energetically unfavorable conformation of HA with acetamido and carboxylate groups on opposing sides of the polymer. (**d)** Position of the gating loop in UDP-bound Xl-HAS-1. (**e)** Gating loop in Rs-BcsA (PDB: 4P00). (**f**) Gating loop in *Candida albicans* CHS-2 (PDB: 7STL). The gating loop is shown in deepteal. Numbers indicate gating loop residue ranges. Conserved residues are indicated in bold and underlined font.

**Figure S7.**
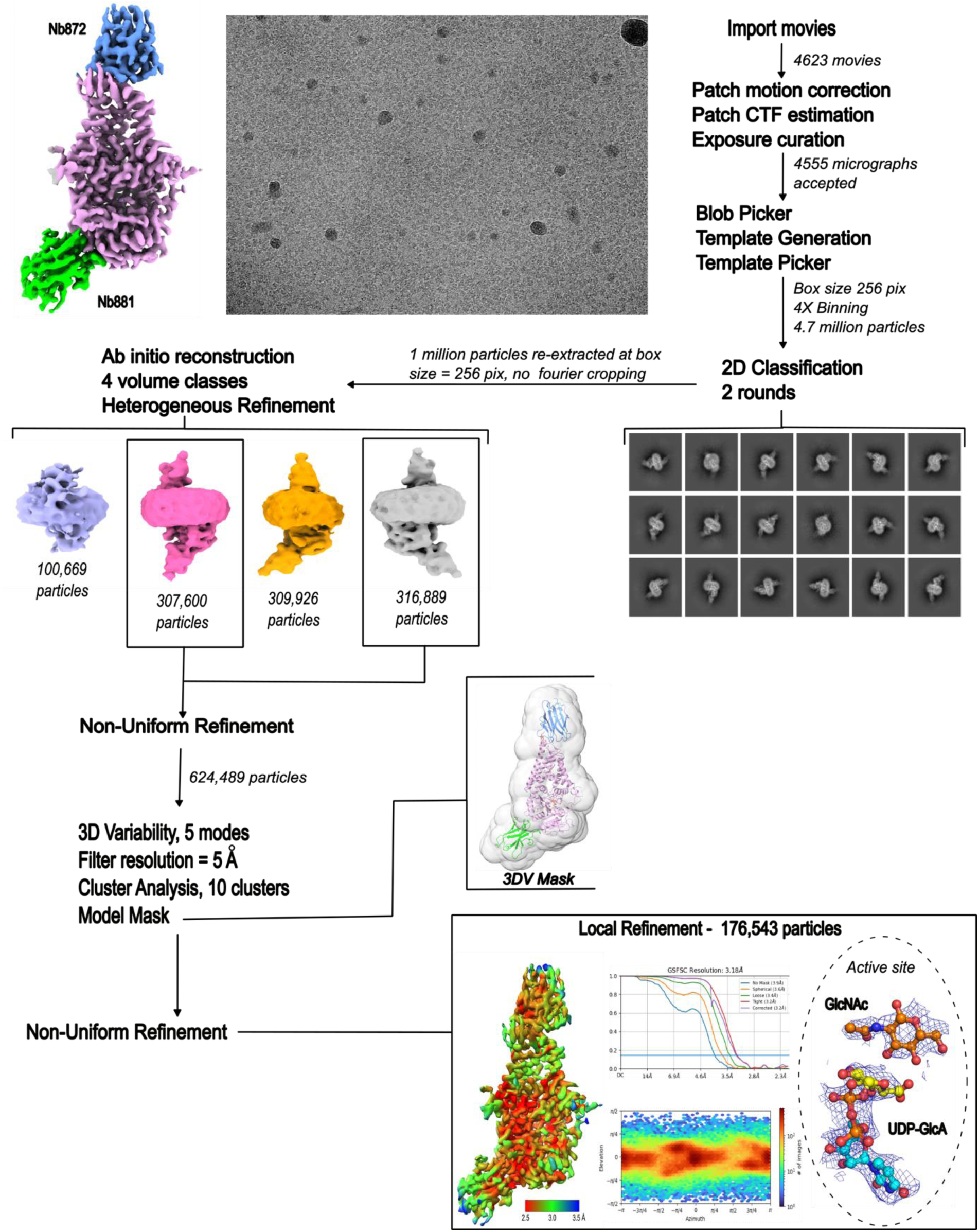
Cryo EM data processing for UDP-GlcA-bound Cv-HAS. CryoEM workflow and map quality for GlcNAc-primed, UDP-GlcA-bound Cv-HAS. UDP-GA and GlcNAc densities contoured at σ=4 r.m.s.d.

**Figure S8.**
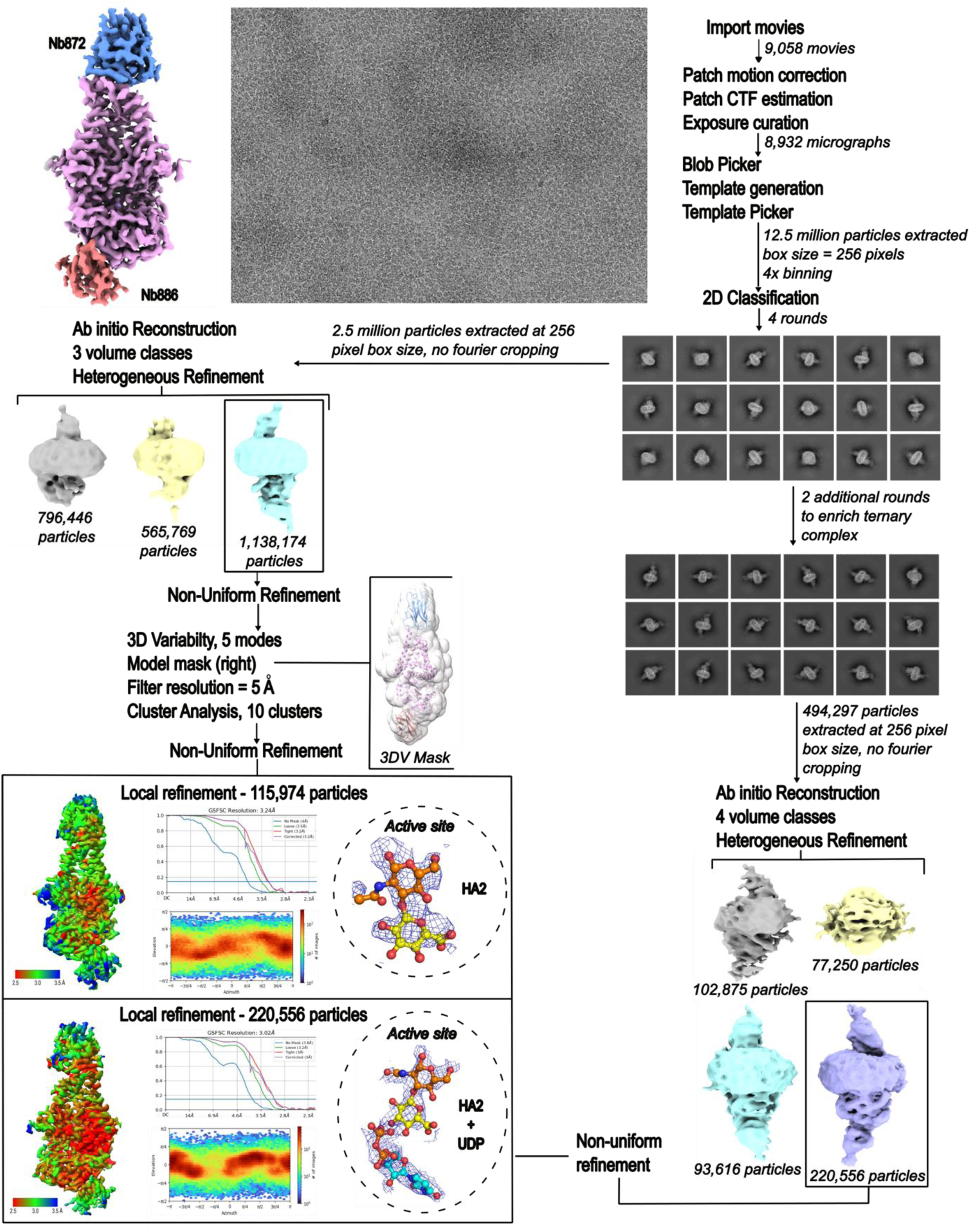
Cryo EM data processing for disaccharide-bound Cv-HAS. CryoEM workflow and map quality for Cv-HAS disaccharide (HA2)-associated Cv-HAS. HA2 and UDP densities contoured at σ=4 r.m.s.d. for both HA2 and HA2 + UDP states.

**Figure S9.**
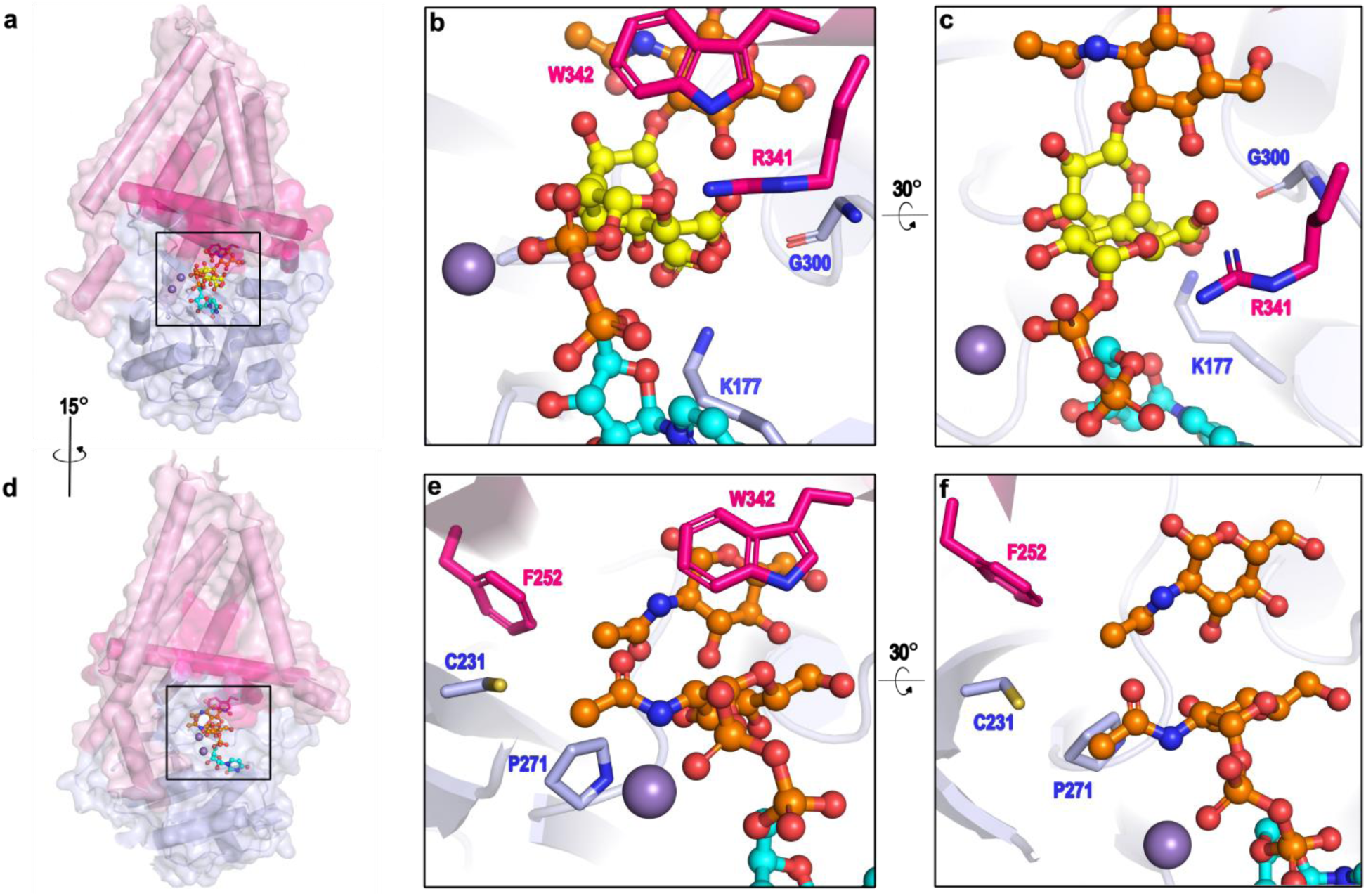
Substrate selectivity due to steric clashes. **(a)**–(**c)**, Superimposition of a disaccharide-bound (HA2) and primed, UDP-GlcA-bound structures for Cv-HAS. (**b**) and (**c**) show shared positioning of UDP-GlcA and HA2 carboxyl moieties in a basic pocket formed by K177, G300 and R341. (**d)–(f**) Overlay of UDP-GlcNAc bound (PDB: 7SP8) and primed, UDP-GlcA bound structures of Cv-HAS. (**e**) and (**f**) show GlcNAc and UDP-GlcNAc acetamido groups occupying the same hydrophobic cavity lined by C231, F252 and P271.

**Table S1.** CryoEM data collection and processing statistics.

| Protein State | XI-HAS-1 |  |  | Cv-HAS |  |  |
| --- | --- | --- | --- | --- | --- | --- |
|  | apo | HA-bound | UDP-bound | Primed + UDP-GlcA | HA2-bound | HA2 + UDP |
| Data collection |  |  |  |  |  |  |
| Microscope | Titan Krios |  |  |  |  |  |
| Camera | K3, GIF, 10eV slit |  |  |  |  |  |
| Magnification | 81,000x |  |  |  |  |  |
| Voltage (kV) | 300 |  |  |  |  |  |
| Dose (e<sup>-</sup>/Å<sup>2</sup>) | 50 |  |  |  |  |  |
| Defocus range (μm) | -2.0 to -1.2 |  |  |  |  |  |
| Pixel size | 1.08 |  |  |  |  |  |
| Data processing |  |  |  |  |  |  |
| Symmetry | C1 | C1 | C1 | C1 | C1 | C1 |
| Particles extracted # | 6,543,217 | 11,732,347 | 14,227,871 | 8,797,858 | 12,554,877 | 12,554,877 |
| Final particles # | 305,106 | 171,641 | 76,968 | 176,543 | 115,974 | 220,556 |
| FSC threshold | 0.143 | 0.143 | 0.143 | 0.143 | 0.143 | 0.143 |
| Map sharpening | 98.4 | 94.5 | 85.9 | 133.0 | 109.1 | 108.6 |
| Model refinement |  |  |  |  |  |  |
| Initial model used | AF2 | XI-HAS-1 apo | XI-HAS-1 apo | Cv-HAS Apo | Cv-HAS Apo | Cv-HAS Apo |
| Model resolution | 3.2 | 3.2 | 3.4 | 3.2 | 3.2 | 3.0 |
| Non-hydrogen atoms | 5993 | 6112 | 6138 | 6089 | 6080 | 6086 |
| Protein residues | 746 | 746 | 762 | 742 | 746 | 743 |
| Ligands | 0 | 9 | 2 | 6 | 6 | 6 |
| B factors |  |  |  |  |  |  |
| Protein | 60.04 | 60.68 | 50.09 | 71.02 | 106.71 | 58.89 |
| Ligand | - | 76.67 | 60.44 | 79.90 | 111.91 | 57.39 |
| Rms deviations |  |  |  |  |  |  |
| Bond lengths | 0.002 | 0.002 | 0.002 | 0.010 | 0.005 | 0.004 |
| Bond angles | 0.461 | 0.462 | 0.505 | 0.935 | 0.997 | 1.024 |
| Validation |  |  |  |  |  |  |
| MolProbity score | 1.56 | 1.57 | 1.57 | 1.20 | 1.66 | 1.52 |
| Clashscore | 5.38 | 6.12 | 6.24 | 3.16 | 6.29 | 4.38 |
| Rotamer outliers (%) | 0.00 | 0.00 | 0.00 | 0.16 | 1.25 | 0.00 |
| Ramachandran plot |  |  |  |  |  |  |
| Favored | 96.06 | 96.47 | 96.55 | 97.54 | 96.34 | 95.65 |
| Allowed | 3.94 | 3.53 | 3.45 | 2.46 | 3.66 | 4.22 |
| Outliers | 0.00 | 0.00 | 0.00 | 0.00 | 0.00 | 0.14 |

